# Axis-specific neuro-musculoskeletal control of body resonance and avoidance behavior during whole-body vibration in mice

**DOI:** 10.1101/2025.11.12.687975

**Authors:** Mamiko Suzuki, Masahiro Saito, Takuya Ueda, Hiroyoshi Inaba, Kou Sasaki, Hirokazu Hirai, Nobutake Hosoi

**Author notes:** Correspondence should be addressed to: Hirokazu Hirai, or Nobutake Hosoi (lead contact),.

## Abstract

Animals must appropriately regulate their responses to external mechanical oscillations such as whole-body (WB) vibration to ensure postural stability. However, knowledge of mouse responses to WB vibration remains limited. Here, we used high-speed video-based quantitative analyses to characterize body-part movements in awake and anesthetized mice under vertical, longitudinal, and lateral WB vibration. Awake mice exhibited lower resonance frequencies and smaller displacement amplitudes than anesthetized mice during vertical and lateral WB vibration, but not longitudinal vibration, indicating that the neuro-musculoskeletal (NMS) system contributes to postural stability in a vibration axis-dependent manner. Vibration modeling suggests that the NMS system acts as an active vibration absorber by reducing effective stiffness and dynamically adjusting effective damping, analogous to active vehicle suspensions. Axis-specific vibration control may reflect an evolutionary balance between postural stability and locomotor efficiency in quadrupeds, paralleling the design principles of rear-wheel-drive vehicles. Behavioral preference tests revealed that mice selectively avoid vertical WB vibration at specific frequencies, but neither lateral nor longitudinal WB vibration. These findings reveal an axis-specific NMS control principle analogous to vehicle design, linking postural regulation, locomotor efficiency and avoidance behavior, and advancing our understanding of the biomechanics and behavior of quadrupeds exposed to WB vibration.

## Introduction

Animals are exposed to various kinds of mechanical vibration, which can be produced by external factors such as interactions with other animals and environmental sources (e.g., vibration during pup transportation by mother animals, ground vibration caused by approaching predators), and internal factors such as self-movement (e.g., head oscillations during running). To maintain stable posture and generate appropriate movements, body parts must respond to such vibratory disturbances in a regulated manner ^1,2^. In humans, whole-body (WB) vibration has been extensively investigated because of its effects on physiological responses, biodynamics, health, and discomfort^3–6^.

By contrast, much less is known about how non-human animals respond to WB vibration ^7,8^. Previous studies in rodents have often measured resonance properties in anesthetized or euthanized animals ^9,10^. This was due to the difficulty of maintaining animals in a fixed position without stress during vibration. Although such approaches yielded valuable basic data on body transmissibility, anesthetized (or euthanized) animals inherently lack active neural and muscular control, thereby overlooking the essential role of the neuro-musculoskeletal (NMS) system in responding to vibration and stabilizing posture ^1,2,11^. Consequently, the dynamics of WB vibration responses in awake animals remain poorly understood. This has been a critical limitation, because the NMS system may actively contribute to vibration attenuation as a spring-damper system ^3–5^ and precise biomechanical understanding of body transmissibility during WB vibration requires data from awake animals in which the NMS system is fully active. In addition, previous studies in animals were focused mainly on vertical WB vibration, neglecting lateral and longitudinal vibration, and measurement points of body transmissibility were very limited (usually a single body point) ^9,10^, due to technical difficulties. These limitations have yet to be fully addressed.

Mice represent a powerful animal model for addressing these issues. As quadrupeds, their postural and locomotor strategies can be compared with those of other mammals, while their small size and pharmacological and behavioral tractability enable precise experimental manipulation and diverse measurements with many experimental toolkits available. Moreover, recent advances in high-speed image acquisition and image processing techniques enable simultaneous monitoring of movements at multiple body parts of a mouse during WB vibration.

Here, we investigated the contribution of the NMS system to WB vibration responses along three axes in mice. Using high-speed video-based analysis, we examined multiple body part responses during vertical, longitudinal, and lateral WB vibration in awake and anesthetized mice. Comparison of these data revealed axis-specific contributions of the NMS system during WB vibration, which is reminiscent of vibration control strategies in rear-wheel drive vehicles. To examine vibration preference in mice, we utilized a two-shuttle-box behavioral test and found a novel vertical axis-specific vibration avoidance behavior that occurs independently of body resonance.

## Results

In the present study, both awake and anesthetized mice were placed in custom-made chambers (mouse boxes) and were exposed to whole-body (WB) vibrations along three axes (vertical [up-down], longitudinal [fore-aft], and lateral [left-right]). The waveforms of the vibrations were sinusoidal with varying frequencies (3 to 40 Hz) and a fixed peak-to-peak displacement amplitude of 1 mm, unless otherwise noticed (Figs. 1–5). The relatively small size of the mouse boxes allowed awake mice to maintain a typical hunched resting posture, which was advantageous for obtaining reliable measurements of body part movements in response to WB vibrations under a consistent posture ^12^ (Supplementary Movies 1 and 2). For anesthetized mice, the feet were affixed directly to the vibrating table to maintain a natural hunched posture as much as possible (Supplementary Fig. S1; see Methods). Movements of various body parts and references in response to WB vibrations were measured by tracking predefined points marked on the skin and the mouse box prior to the experiments (Figs. 1A–3A, and Supplementary Figs. S1 and S2; see Methods). The measured displacement amplitudes of the reference points (∼1 mm, consistent with the intended stimulation amplitude) confirmed that the vibration stimuli were well controlled under all conditions, except at the low frequencies of 3–5 Hz (Supplementary Fig. S3; see Methods). The awake state represents a natural physiological state with intact neural control, whereas the anesthetized state corresponds to a state lacking neural control of body posture ^13^. Comparing awake and anesthetized mice enabled us to investigate the functional contribution of neural control to the musculoskeletal system. Therefore, we quantitatively analyzed the parameters of the measured movements in various body parts, such as peak-to-peak displacement amplitude and phase difference (Supplementary Fig. S2) relative to the reference movement (i.e., input vibration) to examine differences between the awake and anesthetized states. The frequency at which vibration occurs most readily and is amplified in an object, is referred to as the resonance frequency ^8^. We also analyzed the resonance frequencies of multiple body parts in both awake and anesthetized mice during three-axis WB vibration (vertical, longitudinal, and lateral) (Figs. 1–3).

**Figure 1.**
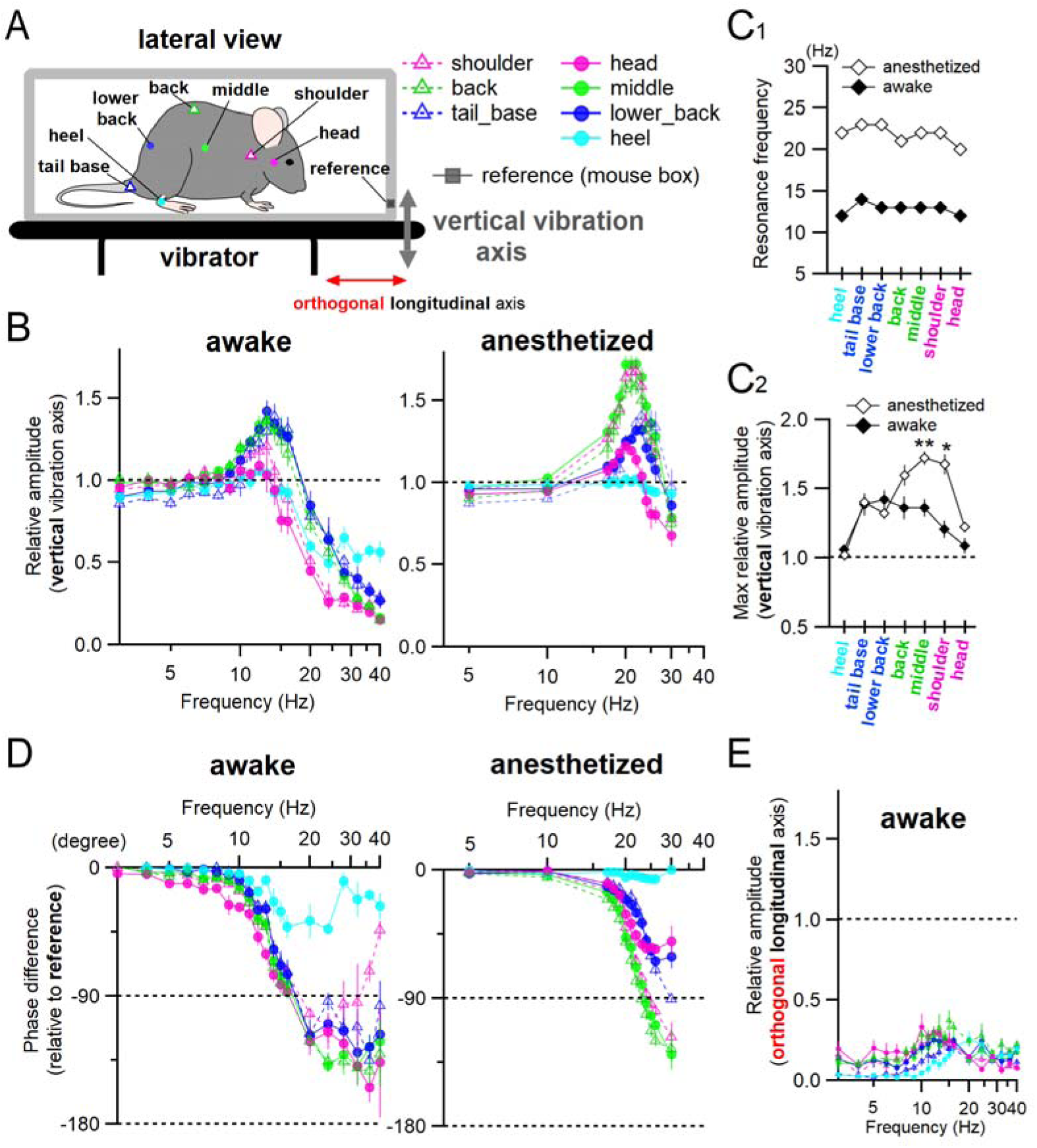
Differential movements of mouse body parts in response to vertical WB vibration in awake and anesthetized mice. (A) Illustration of the experimental setup and the body part locations (tracking points) used to measure displacements during vertical WB vibration (sinusoidal waveform, 1-mm peak-to-peak amplitude at variable frequencies). Color-coded symbols indicate the corresponding mouse body parts analyzed in the other panels (B, D, E). A mouse was placed in a small transparent rectangular acrylic chamber (‘mouse box’) affixed to a vibrator. Body movements were recorded from the side with a digital camera at 1080p (Full HD) resolution and 240 fps during vibration exposure (see Supplementary Video 1). (B) Relative displacement amplitudes of each body part, as depicted in (A), during vertical WB vibration. Measured displacement at each body part was normalized to that of the reference (mouse box), and the resulting relative amplitudes were averaged in awake (left) and anesthetized (right) mice (see Methods). (C_1_, C_2_) Resonance frequency (C_1_; defined as the frequency at which the relative displacement amplitude was maximum in the averaged frequency response profiles shown in (B)), and corresponding maximum relative displacement amplitudes (C_2_) of each body part in awake (filled symbols; n = 6–9) and anesthetized (open symbols; n = 4–10) mice (Mann-Whitney tests between awake and anesthetized mice, \*\**P* < 0.01, \**P* <0.05; see also Supplementary Fig. S4A). Resonance frequencies were lower in all body parts of awake mice compared to anesthetized mice. Notably, the maximum relative displacement amplitudes of anterior body parts (‘head’, ‘shoulder’, and ‘middle’) were reduced in awake mice, whereas those of posterior parts (‘lower back’, ‘tail base’, and ‘heel’) were comparable between the two groups. (D) Mean phase difference of periodic displacement in each body part relative to the reference point in awake (left) and anesthetized (right) mice. Phase difference was first measured as the time lag between the displacement peak of the reference and the temporally adjacent displacement peak of each body part. This time lag was then converted to a phase angle (in degrees), with negative values indicating a phase delay, based on the cycle duration of the vibration frequency (see Methods). (E) Mean relative displacement amplitudes of each body part measured along the longitudinal (fore-aft) axis, which is orthogonal to the vertical vibration axis (i.e., stimulus axis), in awake mice. Displacements along the fore-aft axis were normalized to the vertical displacement amplitude of the reference point (i.e., stimulus input amplitude). Error bars represent the standard error of the mean (SEM) in this and subsequent figures. In (B), (C_2_), and (E), horizontal broken lines indicate the reference amplitude level (equal to the vibration stimulus). In (D), broken lines at -90 and -180 degrees indicate a quarter-cycle and a half-cycle phase delay, respectively.

**Figure 2.**
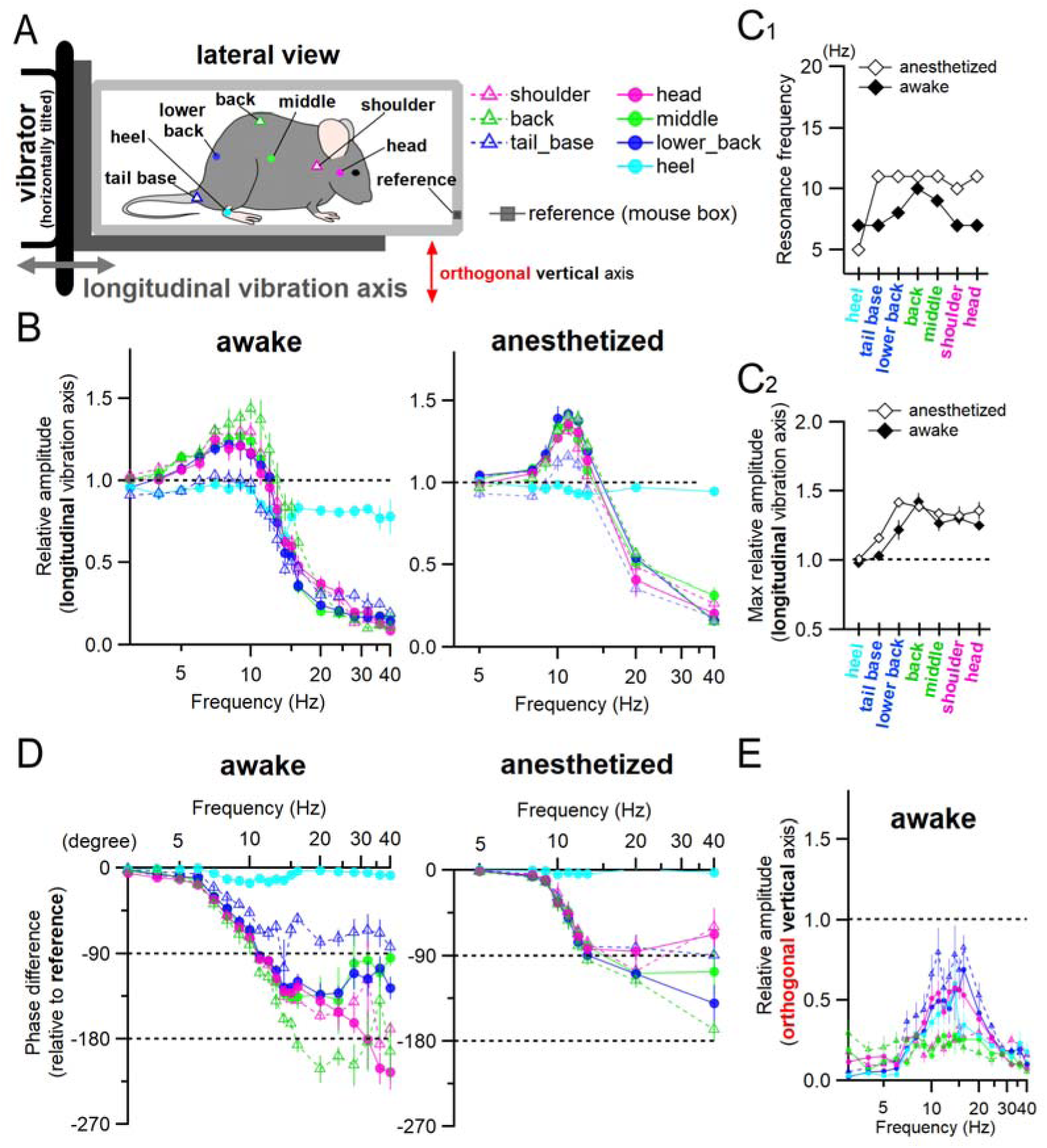
Minor difference in the movements of mouse body parts in response to longitudinal (i.e., fore-aft) WB vibration in awake and anesthetized mice. (A) The experimental setup and measurement locations (tracking points) for mouse body movements were similar to those of Figure 1, except that the vibrator was tilted horizontally. The rear plane of the mouse box was attached to the vibrator to apply longitudinal WB vibration (sinusoidal, 1 mm peak-to-peak amplitude with variable frequency) to mice. (B–E) Data are presented in the same way as in Figure 1. Notably, the reduction in resonance frequency at the body parts of awake mice was minor, especially in the medial parts of the body (the ‘middle’ and ‘back’) and the ‘heel’ (C_1_). The difference in maximum relative displacement between awake (n = 5–6) and anesthetized (n = 6–10) mice was also minor across body parts, except for the ‘lower back’, and ‘tail base’ (C_2_). During longitudinal WB vibration in awake mice, pitching of the mouse body was observed, with pronounced movements along the orthogonal vertical axis in the anterior and posterior parts of the body (E).

**Figure 3.**
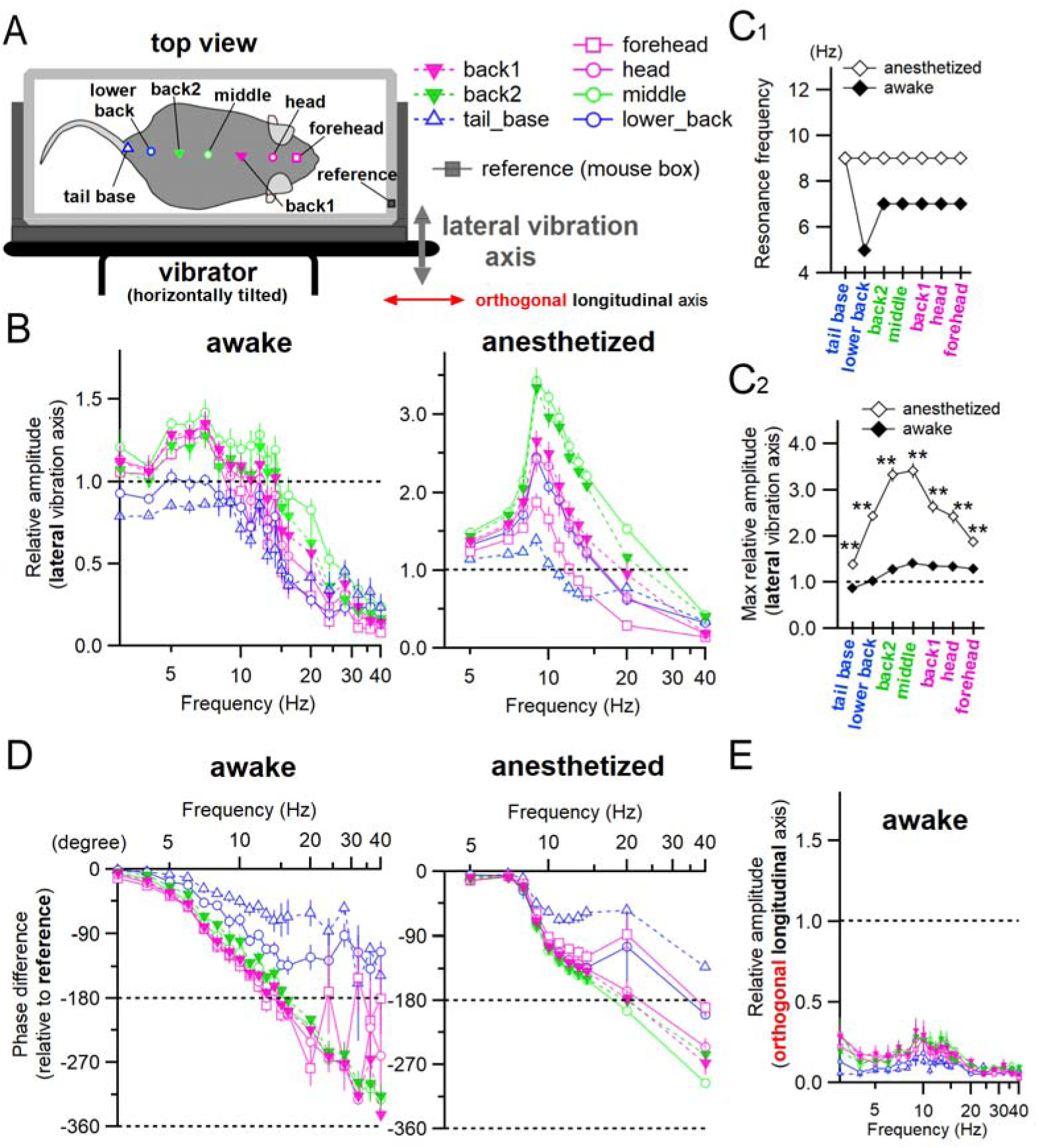
Large difference in the movements of mouse body parts in response to lateral WB vibration between awake and anesthetized mice. (A) The experimental setup was similar to that shown in Figure 1, except that the side plane of the mouse box was affixed to the horizontally tilted vibrator, allowing the application of lateral WB vibration (sinusoidal, 1 mm peak-to-peak amplitude with variable frequency) to mice. Moreover, tracking points for mouse body movements were placed along the midline of the body, and movements were recorded from above. (B–E) Data are presented in the same way as in Figure 1. As with vertical WB vibrations (Fig. 1), resonance frequencies in most body parts were reduced in awake mice compared to anesthetized mice, except for the ‘tail base’ (C_1_). The difference in maximum relative displacement between awake and anesthetized mice was markedly larger than that observed in the other axial WB vibrations (C_2_; Mann-Whitney U-test, \*\**P* < 0.01, comparison between awake [n = 6–7] and anesthetized [n = 6] groups). Movements of the body parts along the orthogonal longitudinal (i.e., fore-aft) axis were negligible during lateral WB vibrations (E).

**Figure 4.**
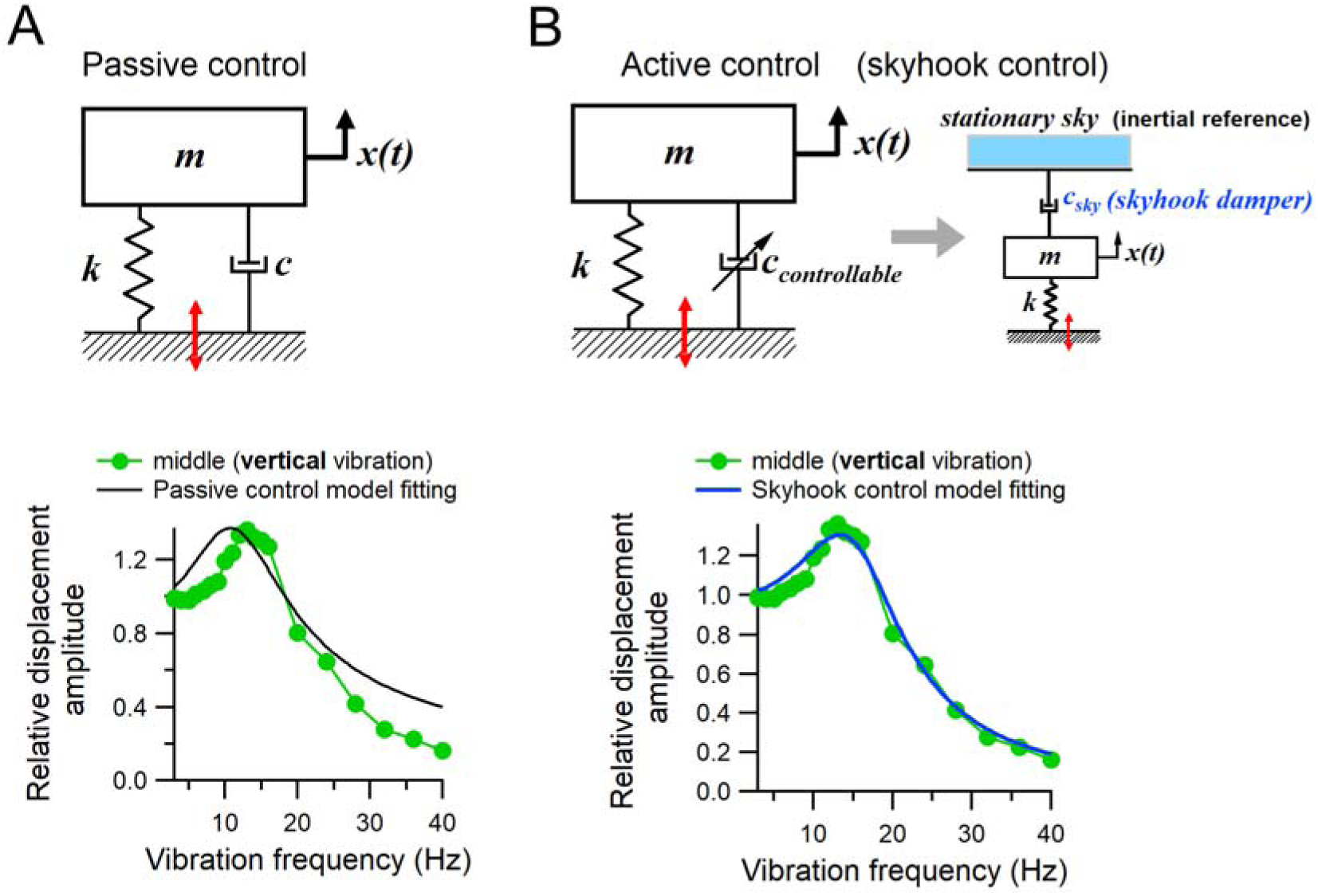
An active control model (skyhook damper control) fits the displacement of the mouse body part ‘middle’ in response to vertical vibration better than a passive control model. (A) and (B) show the two vibration control models, passive control and active control (upper panels), and their corresponding model-based fits to the same experimental data (lower panels), respectively. The measured displacement of the ‘middle’ body part in response to vertical whole-body vibration (Fig. 1) was normalized to the reference amplitude at each frequency (i.e., amplitude of the basal vibration stimulus) and used for model fitting (green filled circles), because the ‘middle’ is approximated as the center of the total lumped mouse body mass. Both models are based on a single-degree-of-freedom vibration system consisting of a lumped mass (*m*) connected in parallel to a spring (stiffness, *k*) and a damper (damping coefficient, *c*). In the upper panels, *x(t)* represents the displacement of the mass at time *t*, and red double-headed arrows indicate the basal vertical vibration input. The mass was fixed at 22.0 g (0.022 kg, corresponding to the typical body weight of an adult mouse), while *k* and *c* (or *c_sky_*) were treated as free parameters for model fitting (see Methods). In the active control model (B), the damper was assumed to be feedback-controlled (i.e., a controllable damping coefficient, *c_controllable_*) and a simple skyhook control scheme was adopted (B, upper panel). In this model, the controlled damping coefficient, *c_sky_* is regulated based on the absolute velocity of the mass. The passive control model yielded estimates of *c* = 2.071 and *k* = 144.7 (A, lower panel, black line), while the skyhook control model provided improved fitting with *c_sky_* = 1.938 and *k* = 239.8 (B, lower panel, blue line).

**Figure 5.**
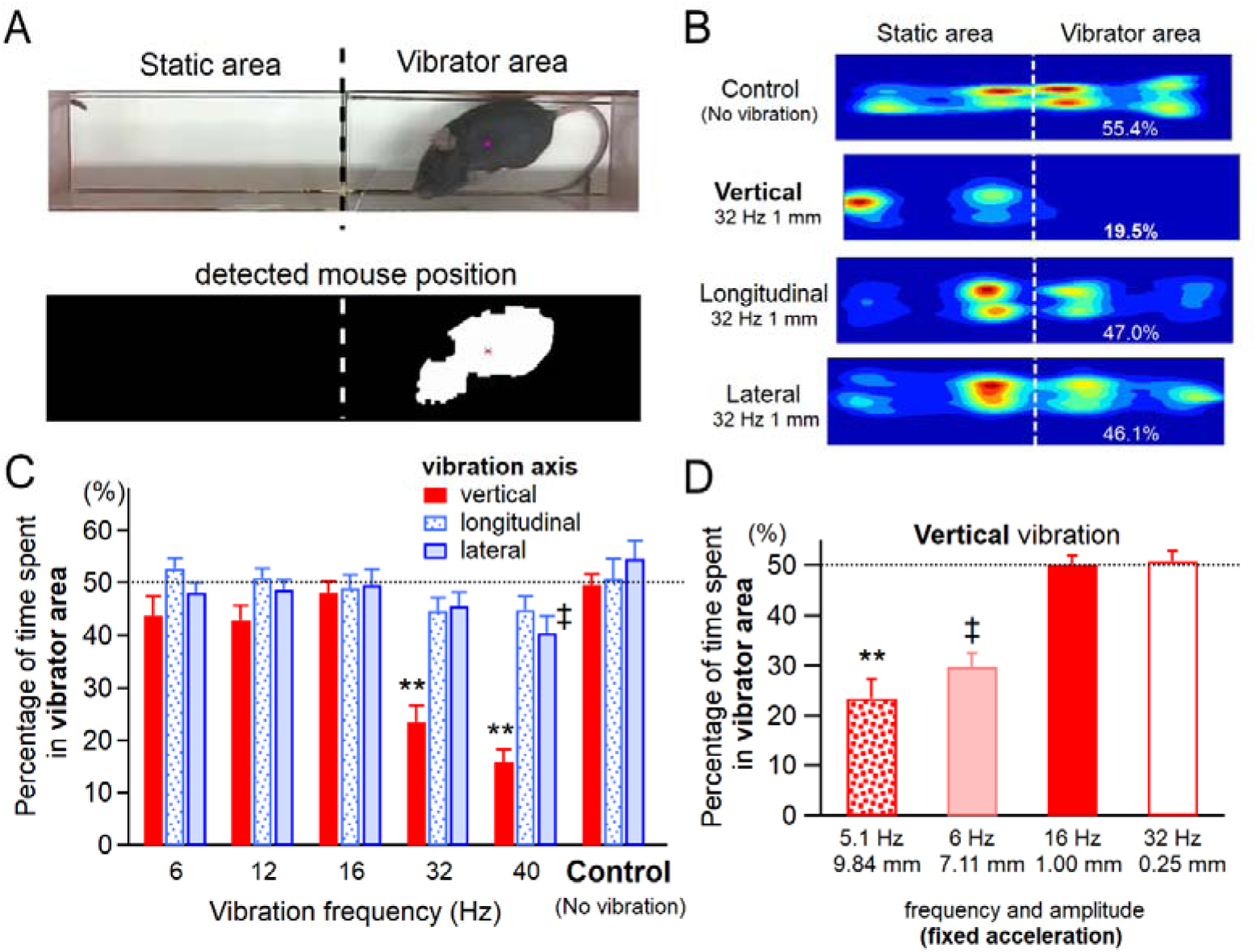
Mice selectively avoid vertical vibration, but not longitudinal or lateral one, depending on frequency and amplitude. (A) Two mouse chambers were placed with their open ends facing each other and separated by a 2-mm gap. One chamber was attached to the vibrator (upper panel, vibrator area), while the other was isolated from vibration (upper panel, static area). Mouse positions were recorded with a digital video camera for 30 minutes to assess approach-avoidance behavior in response to various vibration conditions. The upper panel shows a cropped representative video frame, and the lower panel shows the estimated mouse position (indicated by a red asterisk), determined as the centroid of the mouse body (i.e. a chunk of white pixels) using a color-detection image processing method (see Methods). Broken lines indicate the boundaries between the static and the vibrator areas. (B) Heatmaps showing the mouse’s positional probability for 30 minutes in response to three-axis vibrations. Warmer colors indicate higher probability of presence at each location. Broken lines represent the boundaries between the static and the vibrator areas. White numbers indicate the percentage of time spent in the vibrator area. (C) Mean percentages of time spent in the vibrator area over a 30-min test period, plotted against vibration frequency (1-mm peak-to-peak amplitude) for three vibration axes: vertical (n = 18–21 mice), longitudinal and lateral (n = 18 each). Control data were obtained under no-vibration conditions (vertical, n = 21; longitudinal and lateral, n = 9 each). Dunn’s multiple comparison test, \*\**P* < 0.0001, ‡*P* < 0.005 compared with the corresponding control for each vibration condition. Mice specifically avoid vertical vibrations at frequencies above 30 Hz. (D) Mice also avoided vertical vibrations of larger amplitude at frequencies below 16 Hz, even under a constant peak acceleration of 5.0532 m/s^2^, equivalent to that generated by a vibration stimulus with a 1.00 mm peak-to-peak displacement at 16 Hz (n = 15 mice in each condition; Dunn’s multiple comparison test, \*\**P* < 0.0001, ‡*P* < 0.005 compared to 16 Hz, 1.00 mm or 32 Hz, 0.25 mm). The dotted lines in (C) and (D) indicate chance level.

### Body part movements during vertical WB vibration

Figure 1A depicts the experimental setting and the body part locations used for displacement measurements during vertical WB vibration, with the symbol code also applied to Figure 1B, D, E. Figure 1B shows pooled data of the peak-to-peak displacement amplitudes in awake and anesthetized mice. In awake mice, the relative displacement amplitudes of the middle-to-posterior body parts (‘middle’, ‘back’, ‘lower back’, and ‘tail base’) along the vibration axis were noticeably enhanced within the frequency range of 10–16 Hz (Fig. 1B, left panel; green and blue symbols compared with the black dashed line), indicative of resonance. In contrast, anterior body parts (‘shoulder’ and ‘head’; Fig. 1B, left panel, magenta symbols) exhibited minimal or no enhancement of the displacement amplitude in this frequency range, and damping effects emerged at lower frequencies than the other regions except ‘heel’. Notably, the relative displacement of the ‘head’ remained constant and close to the input vibration amplitude (the black dashed line) at less than 15 Hz, indicating no clear resonance. At frequencies below 10 Hz, displacement amplitudes of all body parts were similar to the reference (i.e., relative amplitude close to 1), although the amplitudes of lower body parts located near the basal vibrating surface tended to be smaller. In anesthetized mice (Fig. 1B, right), the relative displacement amplitudes were substantially larger than those in awake mice across all body parts except for the ‘lower back’, ‘tail base’, and ‘heel’, the three regions located closest to the vibrating surface. This suggests reduced damping (i.e., less suppression of vibration) at most body parts in the absence of active neuromuscular control.

Figure 1C_1_ shows the resonance frequencies, defined as the frequencies at which the relative displacement amplitude reached its maximum for each body part, based on the averaged frequency–amplitude profiles shown in Figure 1B. In all body parts, awake mice exhibited substantially lower resonance frequencies than anesthetized mice. This difference remained consistent and statistically significant even when resonance frequencies were measured from individual frequency-amplitude profiles for each mouse and then averaged by body part (Supplementary Fig. S4A, left; anesthetized mice [n = 6] vs. awake mice [n = 5–7], *P* < 0.05). In the analysis of maximal relative displacement amplitude, awake mice showed significantly smaller amplitudes at anterior-to-mid body regions, specifically the ‘shoulder’ and ‘middle,’ than anesthetized mice, regardless of whether the data were derived from group-averaged (Fig. 1C_2_) or individual profiles (Supplementary Fig. S4A, right).

At frequencies of 20 Hz or higher, awake mice exhibited a profound decrease in displacement amplitudes (i.e. damping effect) across all the body parts, in clear contrast to anesthetized mice (Fig. 1B). Notably, in awake mice, the anterior body parts (‘head’ and ‘shoulder’) and the ‘heel’ began to exhibit a damping effect (i.e., relative amplitudes below 1.0) at approximately 15 Hz, which was lower than the corresponding frequency observed in the posterior body parts (‘middle’, ‘back’, ‘lower back’, and ‘tail base’; approximately 20 Hz). In contrast, anesthetized mice showed only minor damping effects starting around 25 Hz, with a slight decrease in amplitude observed across all body parts at 30 Hz (Fig. 1B, right). At frequencies above 30 Hz, quantification of displacement amplitude and phase difference in anesthetized mice became unreliable due to highly irregular waveforms (e.g., double-peak complex waveforms at the ‘head’ in Supplementary Fig. S5A, B and the ‘lower back’ in Supplementary Fig. S5A), considerable inter-animal variability, and deviation from the simple sinusoidal-like patterns observed in the ‘reference’ (Supplementary Fig. S5). One characteristic complex displacement waveform exhibited a relatively small positive-going hump during the downward phase of displacement, as seen at the ‘tail base’ and ‘shoulder’ in Supplementary Fig. S5A, B. Another distinct pattern, observed at the ‘tail base’, ‘lower back’, and ‘middle’ in Supplementary Fig. S5C, consisted of the fusion of two positive peaks, leading to a broader and larger displacement waveform with an increased apparent cycle time compared to that of the ‘reference’ waveform (i.e., input vibration cycle time). These complex waveform patterns suggest that the body parts of anesthetized mice mechanically behave as coupled oscillators, where interconnected body segments contain two or more oscillators (i.e., a multi-degree-of-freedom system) that interact during vertical WB vibration. Some anesthetized mice exhibited clear amplitude damping at certain body parts (e.g., the ‘shoulder’ in Supplementary Fig. S5A, C; 8 mice), while others showed little damping effect (Supplementary Fig. S5B; 3 mice). Together, these results indicate that the active NMS system in awake mice effectively attenuates body part movements at 20 Hz and higher frequencies, whereas in the absence of neuromuscular control, damping becomes evident only at 30 Hz and higher. Furthermore, the findings suggest that the anterior body parts, particularly the ‘head’ and ‘shoulder,’ are stabilized more effectively by the active NMS system during vertical WB vibration than other regions.

Next, we analyzed the phase difference (i.e., phase delay) of body part movements relative to the ‘reference’ movement during vertical WB vibration (Fig. 1D). Compared to anesthetized mice, awake mice exhibited a greater phase delay (negative phase difference) in all body parts at much lower frequencies. In both awake and anesthetized mice, phase delay emerged around the resonance frequencies of each body part (10–15 Hz in awake mice; 20–25 Hz in anesthetized mice), consistent with predictions from simple vibration theory ^14,15^. Meanwhile, Figure 1E shows that in awake mice, relative displacement along the longitudinal (i.e., fore-aft) axis, orthogonal to the applied vertical WB vibration, was negligible compared to the vertical displacement of the ‘reference’ (i.e., input vertical displacement). These results suggest that mouse body parts may respond to vertical WB vibrations in a manner analogous to a base-excited, forced vibration of a single-degree-of-freedom (SDOF) mass-spring-damper (MSD) system ^3,4,16^. Notably, Figure 1B (left) indicates that the relative amplitude responses of ‘head’ and ‘shoulder,’ which were efficiently damped around their resonance frequencies (10–15 Hz), became smaller at higher frequencies (presumably above √2 times the natural frequency of the system) than those of other, less-damped body parts (e.g., ‘middle’ and ‘lower back’) in awake mice. In other words, there was no adverse (i.e., compromising) effect on high-frequency vibration isolation in the well-damped ‘head’ and ‘shoulder’ compared with the less-damped ‘middle’ and ‘lower back’ (Fig. 1B, left). This response profile in awake mice contradicts the predictions of a simple base-excited forced vibration model ^3,14,17^, and instead favors an active vibration isolation model ^17^ (see Figure 4 and Discussion below).

### Body part movements during longitudinal WB vibration

In awake mice exposed to longitudinal (fore-aft) WB vibration (Fig. 2A), relative fore-aft displacements increased between 5 and 12 Hz, typically exhibiting a single peak in most body parts except the ‘tail base’ and ‘heel’ regions (Fig. 2B, left). In anesthetized mice, relative displacement amplitudes showed similar amplification between 8 and 13 Hz, also with a single peak (Fig. 2B, right). Regarding phase differences in awake mice, the ‘tail base’ and ‘heel’ exhibited relatively small phase shifts, while the ‘back,’ which is the most distant from the vibrating floor, showed a markedly delayed phase response (exceeding -180° at frequencies above 15 Hz) (Fig. 2D, left; see also Methods). Resonance frequencies of body parts in awake mice tended to be slightly lower than those in anesthetized mice (Fig. 2C_1_ and Supplementary Fig. S4B, left). However, no significant differences were observed in the maximum relative displacement amplitudes between the two groups (Fig. 2C_2_ and Supplementary Fig. S4B, right), suggesting little damping effect exerted by the NMS system during longitudinal WB vibration, except for the ‘heel’, which is physically closer to the vibrating floor (Fig. 2B).

Unlike vertical WB vibration, longitudinal (fore-aft) WB vibration produced markedly larger relative displacements along the vertical axis (orthogonal to the stimulus direction) in awake mice within the 10–20 Hz range (Fig. 2A, E). These responses suggest that mouse body movements during longitudinal WB vibration cannot be explained by a single-axis model and instead involve multiple-degree-of-freedom dynamics. In the awake state, with the body supported above the ground by four limbs, longitudinal vibrations may induce rotational, seesaw-like oscillatory vertical movements (pitching) around the body’s center of gravity, located near the ‘middle’ body point (Fig. 2A). This interpretation is supported by Supplementary Video 2 and by the observation that the orthogonal vertical displacements of anterior and posterior body parts, which are farther from the ‘middle’ point, were considerably larger during fore-aft WB vibrations (Fig. 2E; green symbols vs. blue or magenta symbols, except for the ‘shoulder’).

### Body part movements during lateral WB vibration

In contrast to vertical and longitudinal WB vibration, awake mice exposed to lateral WB vibration (Fig. 3A) exhibited complex frequency–displacement response profiles, with two distinct peaks at 7 Hz and 12 Hz in most body parts (Fig. 3B, left). These multi-peaked profiles suggest that body displacements during lateral WB vibration may follow a multiple-degree-of-freedom vibration model in awake mice. Notably, only posterior body parts (i.e., the ‘tail base’ and ‘lower back’) showed displacement amplitudes reduced to or below the amplitude of the applied vibration stimulus (Fig. 3B, left, blue symbols), indicating that these regions are effectively damped and vibration-resistant during lateral WB vibration. Unlike longitudinal WB vibration, orthogonal fore-aft displacements were negligible across all body parts during lateral WB vibration (Fig. 3E), similar to the results observed under vertical WB vibration (Fig. 1E). In anesthetized mice, relative displacement amplitudes were markedly greater than those in awake mice across all body parts, exhibiting single response peaks at 9 Hz (Fig. 3B, right).

Resonance frequencies derived from the group-averaged displacement profiles were lower in awake mice compared to anesthetized mice (Fig. 3C_1_). This trend was confirmed by the analysis of resonance frequencies calculated from individual frequency–displacement profiles (Supplementary Fig. S4C, left). Statistically significant reductions in resonance frequency were observed in the anterior body parts of awake mice compared to anesthetized mice, including the ‘forehead,’ ‘head,’ and ‘back1’ (anesthetized mice [n = 6] vs. awake mice [n = 7], *P* < 0.01).

Analysis of the maximum relative displacement amplitudes along the lateral vibration axis revealed that awake mice exhibited significantly reduced displacements compared to anesthetized mice across all measured body parts, with particularly pronounced suppression observed in the medial regions of the body (‘middle’ and ‘back2’; Fig. 3C_2_; Supplementary Fig. S4C, right). These results suggest that the NMS system suppresses body part movements during lateral WB vibration, functioning as a robust damping and stabilizing mechanism in response to lateral perturbations.

Notably, unlike vertical and longitudinal WB vibration conditions, most body parts in awake mice exhibited phase differences (phase delay) exceeding -180° at frequencies above 15 Hz, with even greater phase delays observed at frequencies above 20 Hz during lateral WB vibration (Fig. 3D, left). At 20 Hz, relative displacement amplitudes remained above 0.27 across all body parts and exceeded 0.5 at the ‘back1’, ‘middle’, and ‘back2’ points (Fig. 3B, left), indicating that phase-difference measurements in this frequency range were reliable. These pronounced phase delays beyond -180° in awake mice may reflect either a passive multiple-degree-of-freedom vibration system or active vibration control mechanisms.

Although Figure 3D shows a gradual increase in phase delay in awake mice and excess of -180° phase difference at around 15 Hz, analysis based on the steady-state response at fixed vibration frequency alone cannot discriminate whether the phase difference is truly delayed beyond -180° or advanced below -180° compared to the reference theoretically. Thus, to investigate how this large phase delay may develop during lateral WB vibration in awake mice, we utilized a frequency-modulated single-sweep vibration stimulus (Supplementary Fig. S6A). The frequency of the stimulus was continuously and gradually changed from 10 Hz to 30 Hz at a slow rate of 0.5 Hz/s (1-mm peak-to-peak displacement amplitude). Moreover, to examine distinctive responses of left and right body parts, symmetrical left and right body parts were included among measurement points, and displacement responses of body parts were measured from the rear of an awake mouse (Supplementary Fig. S7A; see Methods).

Analysis of the data along the lateral vibration axis using the frequency-modulated lateral vibration stimulus suggests that frequency-dependent progressive increase in phase difference continuously develops from around -90° to over -270° in upper body parts (Supplementary Fig. S7C, ‘top’ and ‘upper hip’, magenta and blue), and that characteristics of left and right body parts were similar (Supplementary Fig. S7B and C, filled triangles and solid lines [left] vs. open squares and dotted lines [right]). However, the data analyzed along the vertical axis orthogonal to the stimulus indicate large displacements (reaching over 0.25 mm) of most body parts except for ‘top’ (Supplementary Fig. S7D). This resembles the results of longitudinal (fore-aft) WB vibration experiments (Fig. 2E), implying a similar possibility that lateral WB vibration may induce rotational, seesaw-like vertical oscillations between the left and right sides of the body. To examine this possibility, phase differences along the vertical axis (orthogonal to the stimulus axis) between symmetrical left and right body parts were analyzed (Supplementary Fig. S6B). These results support the above possibility and revealed that vertical body part movements have a constant antiphase (near -180°) relationship between left and right corresponding parts, irrespective of lateral vibration frequencies (Supplementary Fig. S7E).

### Estimation of biomechanical parameters and active damper model fitting

To gain mechanistic insights into how the NMS system contributes to the suppression of body part movements under vertical WB vibration, we made a rough estimation of stiffness *k* (i.e., spring constant) based on an SDOF MSD system subjected to base displacement. We assumed that the measured resonance frequency closely approximates the natural frequency (*f*_n_) of the system. While this assumption may result in a slight underestimation of stiffness *k* due to the presence of damping (Fig. 1 and Table 1), this compromise is unlikely to affect the relative comparison between awake and anesthetized mice. Under this assumption, *k* can be determined by the two parameters: mass (*m*) and resonance frequency, as described in equation [2] in the Methods section and Table 1 ^8,18^. We focused on two body parts, the ‘middle’ and the ‘head’, for *k* estimation. The ‘middle’ is reasonably considered the center of the total mouse body mass and provides a good approximation to the SDOF MSD system with lumped parameters, whereas the ‘head’ is a critical body part since it contains the brain and sensory organs contributing to vision (eyes), audition (ears), and vestibular sensation (vestibules of the ears). The masses of the ‘middle’ and the ‘head’ were assumed to be 22.0 g (0.022 kg) and 2.8 g (0.0028 kg), respectively, based on the typical body weight (BW) of an adult mouse and the mean measured head-to-BW ratio. The estimated stiffness *k* values in awake mice were less than half of those in anesthetized mice (Table 1). This finding suggests that the active NMS system may function to reduce stiffness in response to vertical WB vibration, as interpreted within the framework of a simple base-excited SDOF MSD vibration model. In addition, we made a rough estimation of the damping ratio for the ‘middle’ body part in awake and anesthetized mice. The estimated damping ratio in awake mice was approximately 1.6 times larger than that in anesthetized mice (Table 2), qualitatively suggesting that the NMS system contributes to the suppression of vertical body vibration by increasing the damping ratio of the body.

**Table 1.** Rough estimation of stiffness (spring constants), calculated based on a single-degree-of-freedom system excited by vertical vibration, using fixed body part weights and their corresponding resonance frequencies in awake and anesthetized mice.

| Rough estimation of stiffness, $k$ (N/m) | middle | head |
| --- | --- | --- |
| Awake | 146.8 | 16.1 |
| Anesthetized | 347.4 | 44.6 |

**Table 2.** Rough estimation of the damping ratio for the ‘middle’ body part in awake and anesthetized mice under vertical vibration.

| Condition | Rough estimation of the damping ratio (**Z**) |
| --- | --- |
| Awake | 0.590 |
| Anesthetized | 0.376 |

The above approximation indicates that the NMS system may suppress body part movements through enhanced damping when the NMS system is considered analogous to a base-excited SDOF system with lumped parameters: a mass (*m*) connected to a spring (stiffness, *k*) and a damper (damping coefficient, *c*) ^3,4^. For vibration control, there are two control models in the SDOF-MSD vibration system: a passive control model where a damping property is fixed, and an active control model where damping properties are actively modulated (Fig. 4) ^17^. To examine where the NMS system may behave as passive or active control over external vertical WB vibration, the relative displacement amplitude (i.e., displacement transmissibility) of the mouse body part ‘middle’ in response to vertical WB vibration was fitted with the two control models (see Methods, Fig. 4). The ‘middle’ region was considered the most suitable for model fitting due to the closest region to the mouse’s center of gravity, and the mass was fixed to 22.0 g as described above. As an active control model, we employed the “skyhook” control model where the active control force is applied via feedback depending on the absolute velocity of the mass. The frequency response curve of the relative displacement amplitude in the ‘middle’ was not fitted well to the passive control model, whereas the active control (skyhook control) model explained the measured data reasonably with stiffness k 239.8 (N/m) and *c*_sky_ 1.938 (N•s/m) (Fig. 4). Using these values and equation [2], we estimated more accurate values for the ‘middle’: a damping ratio *Z_sky_* of 0.422 and a natural frequency of 16.62 Hz. These results indicate that the NMS system in mice may function as the active damping control over vertical vibration like an active suspension of a vehicle.

### Mouse preferences for different WB vibration conditions

The above experiments revealed that mouse body-part responses to whole-body (WB) vibration differ markedly depending on the direction of vibration (vertical, longitudinal, or lateral) and its frequency (Figs. 1–3). These characteristics are summarized in terms of displacement magnitude in Table 3. The differential properties of body-part movements under various vibration conditions may influence mouse behaviors, particularly in terms of mouse preferences for WB vibration stimuli ^19^ and potential concerns about adverse effects of resonant vibrations on mice ^9^. Using a two-alternative shuttle box behavior paradigm (Fig. 5A), we examined mouse preferences for various WB vibration conditions (6–40 Hz) along three axes (vertical, longitudinal, and lateral directions) that were all used in the above experiments (Table 3). Interestingly, no significant preference or dislike was observed for either longitudinal or lateral WB vibration at any tested frequency compared to the corresponding control condition without vibration (Fig. 5B, C). A single exception was noted: mice significantly avoided the lateral vibration at 40 Hz, spending only ∼40% of their time in the vibrated chamber (Fig. 5C). In contrast, vertical WB vibration elicited robust avoidance behavior at 32 Hz and 40 Hz (Fig. 5B, C), indicating a clear axis- and frequency-dependent behavioral response. These results suggest that mice selectively avoid vertical WB vibration at frequencies over 16 Hz. This pattern cannot be attributed to body-part resonance, because the resonance frequencies of body parts in awake mice are approximately 6–16 Hz across all vibration axes (Figs. 1C_1_–3C_1_, Supplementary Fig. S4, and Table 3), a range in which no significant avoidance behavior was detected (Fig. 5C). Therefore, we conclude that resonance itself does not account for vibration avoidance.

**Table 3.** Frequency- and vibration axis-dependent displacement of the ‘middle’ and ‘head’ body parts in awake mice in response to 1-mm vibrations.

| Vibratory excitation<br>direction | Displacement at ‘middle’ |  |  |  |  | Displacement at ‘head’ |  |  |  |  |
| --- | --- | --- | --- | --- | --- | --- | --- | --- | --- | --- |
|  | Vibration frequency (Hz) |  |  |  |  | Vibration frequency (Hz) |  |  |  |  |
|  | 6 | 12 | 16 | 32 | 40 | 6 | 12 | 16 | 32 | 40 |
| Vertical (up-down) | — | ↑↑ | ↑↑ | ↓↓ | ↓↓ | — | ↑ | ↓ | ↓↓ | ↓↓ |
| Longitudinal (fore-aft) | ↑ | — | ↓↓ | ↓↓ | ↓↓ | ↑ | — | ↓ | ↓↓ | ↓↓ |
| Lateral (left-right) | ↑↑ | ↑↑ | — | ↓↓ | ↓↓ | ↑↑ | — | ↓ | ↓↓ | ↓↓ |

Abbreviations —, ↑, ↑↑, ↓, and ↓↓ indicate the relative magnitude of displacement in response to vibration along its excitation axis. The ‘middle’ body part is approximated as the center of mass of the mouse. Specifically, —, ↑, ↑↑, ↓, and ↓↓ represent displacements that are equal to, slightly greater than, much greater than, slightly smaller than, and much smaller than (i.e., strongly damped relative to) the stimulus amplitude (1 mm), respectively.

It appears that the higher the frequency of vertical WB vibration was, the more severely mice avoided the vibration (Fig. 5C). However, the displacement peak-to-peak amplitude of the vertical WB vibration was fixed at 1.0 mm, and the energy of the vibration stimulus (i.e., peak acceleration) increased along with the frequency increment in the experiments above. Thus, we next examined whether displacement amplitude itself can affect mouse preferences for vertical WB vibration. To this end, we adjusted both the displacement amplitude and frequency of the vibration stimuli, with their peak acceleration held at a fixed rate of 5.05 m/s^2^, equivalent to that of a 1-mm vibration stimulus at 16 Hz, which did not induce avoidance (Fig. 5C and 5D). In this configuration (fixed peak acceleration conditions), a trade-off relationship exists between displacement amplitude and frequency (Fig. 5D), which allowed us to dissociate the effects of frequency and displacement amplitude under a fixed mechanical energy level. Under these vibration conditions, mice exhibited significant avoidance of vertical vibration when the peak-to-peak displacement exceeded 7 mm at 5.1 and 6 Hz, which are much below the resonance frequency range (Fig. 1B, awake). In contrast, vibration stimuli with smaller displacements and higher frequencies (e.g., 0.25 mm at 32 Hz) did not elicit avoidance, as preference remained near chance level (Fig. 5D). Collectively, these results indicate that both displacement amplitude and frequency influence the aversive response to vertical WB vibration in mice.

## Discussion

In the present study, we analyzed the movements of various body parts during WB vibrations along three different (vertical, longitudinal, and lateral) axes, and compared the response properties between awake and anesthetized mice. To our knowledge, this is the first report to systematically examine resonance frequencies and phase delays in various body parts in awake and anesthetized mice along the three axes, revealing the differential contribution of the NMS system to mouse postural control along the three vibration axes (Figs. 1–3). We found that awake mice exhibit lower resonance frequencies in body parts than anesthetized mice during WB vibrations with all three axes of excitation (Figs. 1C_1–_3C_1_, Supplementary Fig. S4), suggesting that the NMS system functions to maintain body resonance at lower frequencies. Our results indicate that the NMS system suppresses body vibration by reducing stiffness *k* (Table 1), increasing the damping ratio (Table 2), and actively controlling the damper (skyhook damper, similar to vehicle suspension control) when modeled as an SDOF MSD vibration system (Fig. 4). These dual mechanisms—reduction of stiffness and dynamic adjustment of damping—may enable animals to suppress resonance and maintain postural stability during WB vibration. To our knowledge, the present study is the first to propose the functional and mechanical roles of the NMS system in controlling body vibration from a vibration engineering perspective. Using behavioral preference tests, we also examined the behavioral significance of three-axis WB vibrations and found that mice selectively avoid vertical WB vibrations in a frequency- and displacement-dependent manner (Fig. 5).

### A downward shift of body resonance frequency and a softening effect by the NMS system

The functional significance of lowering body resonance frequency by the NMS system in mice remains to be elucidated. From the perspective of vibration isolation, a downward shift in resonance frequency leads to reduction in vibration transmission at frequencies above ∼ √2 times the resonance frequency (more precisely, the system’s natural frequency) ^14^. In vertical WB vibrations, anesthetized mice exhibited body resonance frequencies ranging 20-25 Hz, whereas awake mice showed a lower resonance frequency range of approximately 10-15 Hz (Fig. 1C_1_, and Supplementary Fig. S4A, left). This implies that the mouse NMS system might effectively attenuate vertical body vibrations at frequencies above 15 Hz, a range that could be critical to mice due to potential harm to the brain and internal organs, or disturbances to the visual, auditory, and vestibular systems. From a clinical perspective, the reduced contribution of the neuromuscular system in individuals such as infants, older adults, or unconscious patients may result in elevated body resonance frequencies. This highlights the importance of minimizing exposure to potentially harmful high-frequency WB vibrations during medical transport in ambulances or other vehicles. Such vibrations may approach the resonance range of these vulnerable individuals, leading to amplified body movements and potentially greater physiological stress compared to healthy adults with intact neuromuscular control and lower body resonance frequencies.

So far, it remains unclear how the NMS system reduces the stiffness *k* of body parts (i.e., the ‘softening’ effect) (Table 1), thereby lowering the body resonance frequencies (Figs. 1C_1_–3C_1_, and Supplementary Fig. S4, left). Interestingly, similar phenomena have been reported in rhesus monkeys ^20^ and human subjects exposed to higher magnitudes of vibration ^21–24^. Griffin (1990, Chapter 8.4.2.1) suggested that this effect might result from an involuntary loss of muscle tone by vibration, phasic muscle activity excited by vibration, or the thixotropic behavior of muscles ^18^. All of these hypotheses involve muscle activity ^25^, which is consistent with our findings. However, our results appear counterintuitive, because anesthetized mice, in which all skeletal muscles are relaxed and presumably more softened, exhibit effective higher stiffness; whereas awake mice, in which some (but not all) skeletal muscles are contracted and appear more rigid, show lower effective stiffness during WB vibration when the mouse body is regarded as a vibrating system. This intriguing discrepancy might be reconciled with non-linear properties of the complex posture control system including muscles, tendons, bones, joints, ligaments, and the nervous system ^2,18^. Notably, several studies have reported that muscle stiffness is reduced during and after contraction ^26–29^. In addition, muscle spindle tuning can affect muscle stiffness ^30^, and joint/limb stiffness can be regulated by joint receptors ^31^. As another possibility, the skeletal structures which are intrinsically stiffer, might be dominant to support the body in anesthetized mice, so that body transmissibility is mediated mainly by the skeletal system, while fully relaxed muscles behave primarily as passive mass attached to the skeletal structure during WB vibration.

Zeeman et al. ^10^ reported spinal resonance frequencies of 8–10 Hz during longitudinal WB vibration in anesthetized rats, comparable to our results (Fig. 2B, anesthetized; Supplementary Fig. S4B left, anesthetized). In contrast, Rabey et al. ^9^ estimated much higher resonance frequencies (41–60 Hz in mice) during vertical WB vibration in anesthetized or euthanized rodents than those observed in our study. This discrepancy may arise from differences in resonance estimation methods and vibration stimuli. In particular, Rabey et al. ^9^ employed frequency-modulated sweep vibration stimuli at a rapid rate of 30 Hz/s, which may predominantly induce nonstationary transient responses, whereas our study focused on steady-state responses of the mouse body during vertical WB vibration using discrete, single-frequency vibration stimuli (Figs. 1–3) or much slower frequency-modulated sweep vibration stimuli (Supplementary Figs. S6 and S7; 0.5 Hz/s).

### Active control of body vibration by the NMS system in mice

Our model fitting suggests that the NMS system functions as an active vibration control system to suppress body vibration during vertical WB vibration in mice (Fig. 4). This interpretation is also supported by the frequency response profiles of the more strongly damped anterior body parts in awake mice (Fig. 1B, left). Specifically, the damping sufficient to control resonance at ‘shoulder’ and ‘head’ did not compromise higher frequency isolation; that is, the ‘shoulder’ and ‘head’ did not exhibit larger amplitude responses than the less damped body parts (e.g., ‘middle’ and ‘lower back’) above 20 Hz. This response property is characteristic of active vibration isolation, as utilized in automobile suspensions ^17^. In contrast, the seated human body exposed to vertical vibration has been well described by passive control models (Fig. 4A, upper panel) ^21,32^. This discrepancy might reflect fundamental biomechanical differences: humans are bipedal with an upright rostrocaudal axis, whereas mice are quadrupedal with a prone rostrocaudal axis. Thus, it might be reasonable that mice with four limbs employ a vibration control strategy analogous to four-wheeled vehicles.

Using the fitted parameters (*k* and *c_sky_*) and equation [2] of the active skyhook control model (Fig. 4), the damping ratio (also referred to as the damping factor), *Z_sky_* of the ‘middle’ mouse body (the presumable body’s center of mass) was estimated to be 0.422. This value is comparable to those reported in the Rhesus monkey (∼0.46) ^20^, the seated human body (0.475) ^21^, and walking humans (∼0.3) ^33^ under vertical WB vibration. These findings suggest that the damping ratio of animal bodies in response to vertical vibration may be evolutionarily conserved across species, although it can vary depending on the ethological context of each species. Such conservation might provide biomechanical advantages by optimizing a trade-off relationship between stability against external perturbations (e.g., vibration) and mobility efficiency (i.e., controlled instability with energy cost) during locomotion.

Biological mechanisms underlying active skyhook vibration control in mice remain to be elucidated. In the skyhook damper system, measurement and use of the absolute mass velocity are required; however, in practice, it is not possible to connect a physical damper from a mass (i.e., the mouse body) to an inertial reference ^34^. In mice, head movement-corrected stable visual images together with vestibular signals encoding body acceleration might be exploited to approximate absolute velocity of the body. Group Ia muscle spindle afferents act as muscle velocity sensors ^31,35^, and thus muscle spindle-mediated velocity-dependent feedback control of muscle activity may provide damping of body vibration ^36^, given that damping force is proportional to the velocity of the vibrating body ^14^. Moreover, group Ia muscle spindle afferents and some Golgi tendon organs can respond to vibration in tendons and muscles ^31,35^. Skin, fascia, joint capsules, and ligaments are also innervated by mechanoreceptors that are sensitive to vibration ^35^. Neural processing of proprioceptive information from these sources and the resulting mechanical output could implement active skyhook-like control of body vibration in mice.

### Vibration axis-specific response properties of mouse body parts and its functional implications

In vertical WB vibration, the present study revealed that one of the anterior body parts, the‘head’ is vibration resistant and stabilized most effectively by the active NMS system among mouse body parts (Fig. 1B, awake). A similar effect has also been reported in standing humans ^37,38^. This stabilization may contribute to the stable and accurate acquisition of sensory information, because most sensory organs including eyes (vision), ears (hearing), vestibular apparatus (balance), nose (olfaction), and whiskers (touch), are concentrated in the head where stability is critical for detecting and processing these sensory stimuli in mice. Moreover, head stability may protect the brain from mechanical damage during locomotion which inevitably involves vertical WB vibration.

In anesthetized mice where the NMS system is inhibited, only vertical WB vibration caused complex. coupled oscillation-like displacement patterns of body parts (Supplementary Fig. S5), whereas no such patterns were observed during longitudinal or lateral WB vibration. This phenomenon may result from the effect of gravity along the vertical axis. In awake mice, NMS system-mediated vibration control may suppress such coupled oscillations which could otherwise be injurious to certain body parts, and instead allow the body to respond as a simple SDOF vibration system (Figs. 1 and 4), thereby minimizing mechanical stress among body parts. Limiting the system’s degrees of freedom may also be advantageous by reducing the cost of body control ^39^.

The present study found that the mid-body regions, ‘middle’ and ‘back’ move maximally among the body parts, regardless of vibration axis or the state of the NMS system (active or inactive) (Figs. 1–3 and Supplementary Fig. S4). This phenomenon may reflect that the mid body, including internal organs, lacks rib support, making it more susceptible to vibration.

Curiously, one of the posterior body parts, the ‘lower back’ in awake mice tended to exhibit larger vibration responses than in anesthetized mice lacking the NMS system-mediated control during vertical WB vibration (Fig. 1B, blue circles, 1C_2_; Supplementary Fig. S4A, right, ‘lower back’). This finding is in sharp contrast to our general observation that the NMS system suppresses vibration responses of body parts in awake mice (Figs. 1-3) and was not observed during longitudinal (fore-aft) or lateral WB vibration (Figs. 2B, C_2_, and 3B, C_2_; Supplementary Fig. S4B, C, right). These results suggest that vertical axis-specific response enhancement at the ‘lower back’ might contribute to active postural control against gravity by the NMS system. During vertical WB vibration, the NMS system might functionally couple the posterior body parts (‘lower back’, ‘back’, ‘middle’, and ‘tail base’) to reduce mechanical stress within the posterior body area.

Among the three axes of WB vibration, two distinct response properties were observed only during longitudinal WB vibration in mice. First, vibration responses were largely uniform across body parts, except for lower body regions close to the vibrating floor, in both awake and anesthetized mice (Fig. 2B). This indicates that, unlike in the vertical and lateral axes (Figs 1B and 3B), mouse body parts are equally mobile along the longitudinal axis. Second, displacement responses differed only slightly between awake and anesthetized mice (Fig. 2B, C_2_), suggesting that the NMS system exerts the least control over body posture along the longitudinal (fore-aft) axis. In other words, mouse body parts remain easily movable in the longitudinal direction even when the NMS system is active. Analogous to four-wheeled vehicles, such mobility may be advantageous for quadrupeds like mice, which primarily locomote in the fore-aft direction. In addition, interestingly, mouse stride frequency during wheel running typically ranges from 4 to 9 Hz ^40^, which corresponds well to our observed resonance frequency range during longitudinal WB vibration in mice (Fig. 2B and 2C_1_, awake; Supplementary Fig. S4B, left). This correspondence may also contribute to efficient locomotion in mice.

Regarding the lateral (left-right) axis, it is noteworthy that the posterior body parts of mice are least vibrated among all body parts during lateral WB vibration (Fig. 3B blue symbols, 3C_2_; Supplementary Fig. S4C, right). This phenomenon indicates that the anterior and central body parts remain relatively mobile, while the posterior parts are comparatively stable and immobile along the lateral axis. Such differential stability may play a critical role in efficient control of body direction during locomotion. In mice, the hindlimbs are mainly used to generate propulsion and accelerate the body, whereas the forelimbs can contribute to braking and steering ^41,42^. Thus, posterior stability may facilitate propulsion, whereas anterior mobility may be advantageous for steering. The mouse body may be analogous to a rear-drive four-wheeled vehicle, complementing previously reported left-right locomotor asymmetries ^43^. Taken together, axis-specific modulation of body vibration control may represent an adaptive strategy that balances locomotor efficiency with attenuation of inappropriate body vibration.

### Vibration discomfort in mice: relationships with body transmissibility and vibration axis

Our behavioral results using the shuttle-box paradigm include two types of behaviors: escape, where mice move away from immediate and present aversive stimulus (i.e., “Get me out of this vibration NOW”), and passive avoidance, where mice show preventive behavior against predicted future threats (i.e., “Do not let me get into that vibration”). Our analysis (Fig. 5; percentage of time spent in the vibration area) might primarily quantify the passive avoidance component. Notably, our behavioral results showed no carry-over effect, and the aversive vibration stimuli were not associated with specific chamber locations (i.e., vibration areas) in mice (see Methods). Therefore, our behavioral results likely represent vibration discomfort corresponding to a mild, rather than strong, unpleasant emotional state in mice.

In humans, it has been suggested that vibration discomfort can be related to body transmissibility and resonance phenomena within the body ^3,6^. However, only weak correlations have been found between vertical head motion and discomfort, and psychophysically measured equivalent comfort contours do not mirror the inverse of body transmissibility curves in humans ^3^. Rather, vibration frequency is considered the primary determinant of human vibration discomfort ^44^. This is consistent with our findings that vibration discomfort in mice depends on vertical vibration frequency when displacement amplitude is fixed, irrespective of resonance frequencies at specific body parts (Table 3; Fig. 5C). Pain after WB vibration exposure also has been reported to be frequency dependent and independent of the resonance frequency in rats ^45^. Vibration discomfort is thus a complex phenomenon determined by multiple internal and external factors, and cannot be predicted solely by the response of a single body part or a single vibration variable ^3,44^. Consequently, general models of vibration discomfort remain elusive, and existing models have been developed under specific experimental conditions with distinct constraints in humans ^3^. Notably, when vertical vibration acceleration is held constant, discomfort in mice becomes dependent on displacement amplitude rather than vibration frequency (Fig, 5D), highlighting the dynamic interplay between frequency, displacement amplitude, acceleration, and environmental context of vibration.

The present study demonstrated that vibration discomfort in mice is specific to vertical WB vibration (Fig. 5). Similar vertical axis specific phenomena have been reported in humans, where vertical vibration induced greater discomfort than horizontal vibration (lateral or longitudinal vibration) at frequencies of 4 Hz and above ^44,46^. This indicates that heightened sensitivity to vertical vibration may be evolutionarily preserved in mammals. Such axial (directional) dependency is reminiscent of a human cognitive function, the inversion effect in visual perception, such as Thatcher illusion (face inversion effect: inverted faces reduce sensitivity to notice face distortion) ^47^ and the shape-from-shading inversion effect (convex-concave perception changes, depending on lighting cues based on the human visual system assumption that light comes from above) ^48^. Comparable face inversion effects along the vertical axis have been reported in non-human primates ^49^ and medaka fish ^50^. Because animals on earth are affected by gravity and gravity always acts along the vertical axis, animals may have adapted themselves to this constraint through evolution. Therefore, vertical WB vibration may possess a unique biological significance in eliciting behaviors such as vibration avoidance in mice.

In conclusion, we identified axis-specific features of body posture control and avoidance behavior under three-axis WB vibration in mice. It remains to be clarified how such axial specificity is achieved from biomechanical and neurophysiological perspectives. Interestingly, some parallels were observed between vibration control in mice and control strategies in rear-drive four-wheeled vehicles. Analogies between quadrupeds and vehicles ^43^ may provide a useful framework for future studies on the biomechanics and NMS system-mediated control of posture in mice.

## Methods

### Animals

All procedures for the care and treatment of animals were carried out in accordance with the Japanese Act on the Welfare and Management of Animals and the Guidelines for the Proper Conduct of Animal Experiments issued by the Science Council of Japan. The experimental protocols were approved by the Gunma University Animal Care and Experimentation Committee (approval numbers: 18-019 and 23-018). Adult C57BL/6 mice of both sexes (BW 17.3–25.3 g) were used for body part movement analysis, and male mice (BW 21.8–31.7 g) were used for vibration avoidance (preference) tests. The mice were maintained on a 12:12-hour light-dark cycle (lights on at 8 a.m.) and at a constant temperature of 23±3℃. Rodent laboratory chow (MF, Oriental Yeast Co., Ltd) and water were freely available in their home cages.

### Vibration apparatus and video recording of body-part movements in awake and anesthetized mice

Mice were exposed to sinusoidal base oscillations at frequencies ranging from 3 to 40 Hz with a constant peak-to-peak displacement amplitude of 1 mm in three different axial directions: vertical (up-down: z-axis), longitudinal (fore-aft: y-axis), lateral (left-right: x-axis). We utilized video object tracking to examine the vibration response in various parts of the mouse body (see Methods below; Supplementary Videos 1 and 2). Unless otherwise noted, the displacement amplitude of the vibration stimuli was set to 1 mm, as our preliminary experiments showed that 1-mm vertical vibrations at 3–56 Hz for 1 hour had no effect on the behavioral results of open field tests conducted immediately after exposure to the vibrations (n = 4–10), suggesting that 1-mm vibrations are not harmful to mice. Moreover, with 1-mm vibration stimuli, our video-based tracking method provided sufficient spatial resolution to detect mouse body part movements across a frequency range of at least 3–25 Hz, which covered the resonance frequencies for all three vibration axes in both awake and anesthetized conditions (Figs. 1–3; Supplementary Figs. S2 and S4). An electromagnetic vibratory actuator (m060, IMV CORPORATION, Osaka, Japan) was used to generate vibration. The actuator was controlled by a vibration controller system (K2, IMV Corporation, Osaka, Japan) connected with a signal amplifier (MA1, IMV CORPORATION, Osaka, Japan). The magnitude (acceleration) and frequency of the base vibrations were monitored with an accelerometer (356A32, PCB Piezotronics, NY, USA) placed on the vibrator platform. In experiments using awake mice, no sedatives or anesthesia were administered. Mice were placed in a transparent acrylic chamber (mouse box; 100 mm length × 40 mm width × 40 mm height) with a detachable lid composed of a thin plastic sheet and thick polyurethane foam containing multiple ventilation holes. This chamber size allowed mice to maintain a natural hunched posture continuously without stress. For vertical vibration, the chamber was secured to the actuator’s vibrating table with double-sided adhesive tape or with instant glue applied onto polyimide tape, which was attached to both the table surface and the chamber bottom (Fig. 1A). For longitudinal or lateral vibration, the actuator was rotated 90 degrees (i.e. positioned horizontally), and an L-shaped attachment was screwed onto the vibrator platform. Depending on whether the rear or side panel of the mouse chamber was fixed to the vibrator side of the attachment, this setup enabled longitudinal or lateral vibration, respectively (Figs. 2A and 3A). A sheet of paper-type bedding (Pulmas 3000, Scitex, Kawasaki, Japan) was placed on the chamber floor to absorb urine and feces. In experiments under anesthesia, mice were pretreated with a ketamine (100 mg/kg, body weight) and xylazine (10 mg/kg) mixture dissolved in 0.9% NaCl solution, administered intraperitoneally (100 µL/10 g). To approximate a natural hunched posture, anesthetized mice were positioned with their limbs folded under the body ^9^, and all paws were affixed directly to the vibrating table (for vertical vibration) or to an attached acrylic plate (for longitudinal and lateral vibration) using four strips of double-sided adhesive tape (Supplementary Fig. S1B, C). The positions of the adhesive stripes were fixed and bilaterally symmetric (Supplementary Fig. S1A). Unless otherwise noted, mouse body movements were recorded using the slow-motion video acquisition mode (1080p Full HD resolution at 240 frames per second [fps]) of an iPhone 8 Plus (Apple Inc., Cupertino, CA, USA), positioned laterally (for vertical and longitudinal vibrations; Figs. 1A and 2A, Supplementary Fig. S1B, Supplementary Videos 1 and 2) or dorsally (for lateral vibration; Fig. 3A, Supplementary Fig. S1C) relative to the mouse or the mouse chamber. The recording duration at each frequency was typically 1 min or less (up to a maximum of 5 min), with inter-trial intervals of at least 0.5 min. The order of stimulus frequencies was randomized.

When a frequency-modulated, single-sweep vibration stimulus was used, the waveform was generated using a built-in function of the vibration controller system (K2, IMV Corporation, Osaka, Japan). The stimulus frequency was linearly increased from 10 Hz to 30 Hz at a slow rate of 0.5 Hz/s, with a constant peak-to-peak displacement amplitude of 1 mm (Supplementary Fig. S6A).

### Measurements of displacement and phase differences at various body parts

To track body movements reliably, mouse body hair was shaved and tracking points on several body parts were marked directly on the skin using a whiteout pen prior to video recording (Supplementary Fig. S1B and S1C; Supplementary Videos 1 and 2). In vertical and longitudinal (fore-aft) vibration, measurements were performed at seven points along the lateral side of the body: ‘head’, ‘shoulder’, ‘middle’, ‘back’, ‘lower back’, ‘tail base’, and ‘heel’ (Figs. 1A and 2A). In lateral vibration, body-part movements were measured from the dorsal side at seven regions arranged in rostrocaudal order: ‘forehead’, ‘head’, ‘back1’, ‘middle’, ‘back2’, ‘lower back’, and ‘tail base’ (Fig. 3A). As reference markers for vibration, solid objects such as the ‘mouse box’ affixed to the vibrating floor, were marked with black dots using a black oil-based pen (Figs. 1A-3A; Supplementary Fig. S1B, C).

To focus on the steady-state response to vibration, body part movements were analyzed at least 5 seconds after vibration onset. Because awake mice were able to move freely in the chamber even during vibration, analysis was limited to video segments in which the mouse maintained a stable hunched posture. Tracking of the marked points in the recorded videos was performed using the open-source software Kinovea (version 0.8.26; https://www.kinovea.org), with the marker tool positioned at the center of each marked point (Supplementary Videos 1 and 2). Of the measured coordinates (x and y in pixels) over time (ms), the coordinate axis either parallel or orthogonal to the vibration direction was selected for analysis, depending on the direction of the vibration stimulus. For example, in vertical vibration experiments, y-coordinate values were analyzed as a function of time (Fig. 1B, D), whereas x-coordinate values were analyzed in Fig. 1E. The values in pixels were converted to millimeters based on calibration scales derived from known dimensions of objects such as the ‘mouse box’ and a yellow reference object (Figs. 1A–3A; Supplementary Fig. S1B, C). Tracked data exhibiting an unstable baseline drift due to postural changes were excluded from the analysis.

From the tracked data segments containing at least five stable cycles of vibration response at each body part, peak-to-peak displacement amplitudes (i.e., double amplitudes) and phase differences relative to a reference were analyzed using a custom-made macro in Igor Pro 8.04 software (WaveMetrics, Lake Oswego, OR, USA) by NH (Supplementary Fig. S2). Positive and negative peak positions were detected using the built-in “FindPeak” function in Igor Pro (box size for sliding average: 4–6), and all detections were visually confirmed (Supplementary Fig. S2, cross symbols). Peak-to-peak displacement was calculated as the difference between adjacent positive and negative peaks of the displacement waveform (Supplementary Fig. S2, black double-headed arrows). All displacement amplitudes were normalized to that of the ‘reference’ in each trial. Phase difference was defined as the delay relative to the adjacent ‘reference’ peak. Specifically, the delay was initially calculated as the time difference between the displacement peak (positive or negative) of the ‘reference’ (Supplementary Fig. S2; red vertical dotted lines) and the nearest subsequent peak of the same polarity in the displacement waveform of each body part (Supplementary Fig. S2; black vertical dotted lines). This delay (Supplementary Fig. S2; red arrows) was then converted to an angle (degrees) based on the cycle time of the vibration frequency (with a full cycle time delay corresponding to -360 degrees). For each mouse, averaged values of displacement and phase difference were calculated from at least six successive cycles of the displacement waveform. At higher vibration frequencies, where displacement amplitudes were often greatly reduced (i.e., strongly damped) and approached the detection limit, the waveforms became nearly flat, making peak detection unreliable. This led to large variability in the measured phase differences and compromised the accuracy of vibration-response quantification. Such unreliable data were excluded from further analysis, resulting in smaller sample sizes at higher frequencies. All data are presented as mean ± SEM (awake mice, n = 2–10; anesthetized mice, n = 4–10). Due to technical limitations of the vibrator at low frequencies, vibrations at around 3 Hz could not be precisely controlled. Although the vibrator was set to generate vibrations with 1-mm amplitude at 3 Hz, the actual ‘reference’ displacement was reduced slightly (Supplementary Fig. S3).

To examine the frequency-dependent progressive increment of phase difference and the interrelationship between left and right body parts, we utilized the single-sweep frequency-modulated lateral WB vibration stimulus (see Methods above). Symmetrical left and right posterior body parts were designated as measurement points, and body part movements were recorded from the rear view (Supplementary Fig. S7A). Measurements of stimulus frequencies, displacements and phase delays were described in Supplementary Figure S6 and performed using a custom script of MATLAB (version 9.9.0, R2020b; The MathWorks Inc., Natick, MA, USA) written by M. Suzuki. The results obtained with the frequency-modulated stimulus were largely comparable to those obtained with the fixed frequency stimulus (Supplementary Fig. S7), indicating that the modulation rate was sufficiently slow to allow steady-state responses of the body parts.

### Model fitting

To describe the dynamic response of the mouse body, a simple single-degree-of-freedom (SDOF) vibration model with lumped parameters was employed. This model consists of a single moving mass supported by a spring and damper, excited by base displacement (Fig. 4) ^3,4^. For model fitting, we focused on the displacement response of the ‘middle’ body part in awake mice under vertical vibration, as a simple and reasonable approximation of the whole-body dynamics. The ‘middle’ body part was considered to approximate the center of the total body mass (*m*, in kg). The mass parameter *m* was fixed at 0.022 kg (22.0 g), corresponding to the typical body weight of an adult mouse. In the passive control model of base excitation (i.e., vibration applied at the base), both the spring coefficient (*k*, in N/m) and damping coefficient (*c*, in N·s/m) were assumed to be constant. According to the model, the relative amplitude of the mass displacement with respect to the base (input) displacement (i.e., transmissibility) is given as a function of the stimulus frequency (*f*, in Hz) as follows:

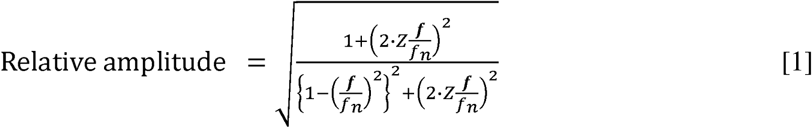

where *f_n_*is the natural frequency, and Z (usually corresponding to ζ) is the damping ratio, defined as:

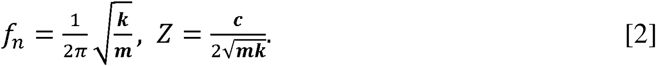

Model fitting was performed to the experimental data using this equation, treating *k* and *c* as free parameters (Fig. 4A). In the second model, an active control approach incorporating a dynamically controllable damper was employed. Specifically, we adopted an ideal skyhook damping control system (Fig. 4B) in which the damping force is regulated via feedback based on the absolute velocity of the mass ^34^. This “skyhook” scheme was chosen for its simplicity in mathematical formulation. In this model, the damping coefficient *c_sky_* serves as a feedback gain proportional to the absolute velocity. In this model, the relative amplitude is expressed as:

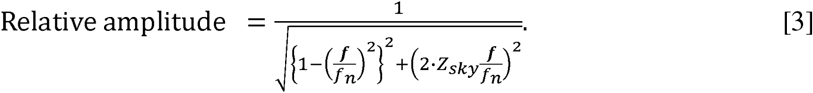

Here, *Z_sky_* is defined in the same way as *Z,* except that the damping coefficient, *c* is replaced with *c_sky_*. The model was fitted to the experimental data using *k* and *c_sky_* as free parameters (Fig. 4B). Model fitting was limited to data from awake mice, as data from anesthetized mice exhibited complex responses suggestive of a multi-degree-of-freedom system at higher frequencies (Supplementary Fig. S5), which made them unsuitable for the simplified models described above. In addition, model fitting was not performed to data from lateral or longitudinal WB vibrations, because these conditions were also complex, indicative of a multi-degree-of-freedom system, and likely violated the assumption of a simple SDOF system.

### Rough approximation of the damping ratio

The damping ratio (Z; corresponding to ζ) was roughly estimated for the body part ‘middle’ in both awake and anesthetized mice from the following equation describing the maximum relative displacement amplitude (i.e., maximum displacement transmissibility *T_max_*):

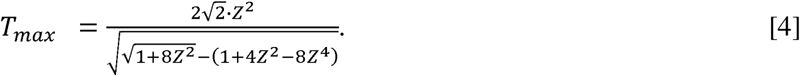

This equation is derived from a passive mass–spring–damper system subjected to harmonic base displacement (Fig. 4A). The damping ratio Z was determined by substituting the experimentally measured peak relative amplitude for *T_max_* with and numerically solving the equation [4] using the ‘fsolve’ function in the SciPy Python library. To avoid dependence on the initial value, the calculation was repeated with initial guesses ranging from 0.1 to 1 in 0.1 increments, and the appropriate converged solution was selected. This approximation should be interpreted as a rough estimate and was used only for qualitative comparison of damping properties between awake and anesthetized mice.

### Preference (avoidance) test for vibration

All animals were habituated to experimenters by daily handling for 5 days prior to behavioral testing. During handling, each mouse was placed on the gloved palm of the experimenters for 10 minutes per day.

To examine vibration preference (or avoidance) in mice, a two-chambered shuttle-box behavioral paradigm was employed (Fig. 5A). The apparatus consisted of two open-ended acrylic rectangular chambers of identical size (130 mm [length] × 45 mm [width] × 45 mm [height]). The top panels were transparent to allow overhead video recording of the mouse, while the remaining panels were opaque grey to reduce exposure to external visual stimuli. The chambers were arranged such that their open ends faced each other, separated by a 2-mm gap, allowing the mouse to shuttle freely between them.

One chamber was affixed to the vibrating table (vibrator area), while the other was placed on an immobile platform of identical height (static area). A sheet-type white bedding (Pulmas3000, Scitex, Kawasaki, Japan) was taped to the floors of both chambers to absorb urine and feces, and to enhance color-based detection of the mouse body (black-colored) during image analysis. Illumination intensity ranged from 245 to 432 lux, and the brightness levels in both chambers was adjusted to the similar level so that the difference between them was less than 10 lux.

Before testing, mice were transported to the testing room and acclimated for 20 minutes in a plastic cage (260 mm [length] × 155 mm [width] × 125 mm [height]) housing three mice per cage. Following acclimation, each mouse was placed in the vibration chamber (Fig. 5A; vibrator area), where it voluntarily entered without signs of distress. The opening between the two chambers was closed with a removable guillotine-style partition, and the vibration stimulus was applied for 1 minute to allow sufficient exposure for preference formation. Subsequently, the partition was removed, and the mouse was allowed to move freely between the two chambers for 30 minutes. All tests were conducted in a quiet room between 9 a.m. and 12 p.m. Each mouse was tested once per day with an interval of at least one day between trials. On the first day, a habituation trial was conducted without vibration to allow mice to become familiar with the shuttle-box environment. From day 2 onward, mice were exposed to vibration stimuli presented in randomized order. The vibration conditions included a fixed peak-to-peak displacement amplitude of 1 mm at different frequencies (6, 12, 16, 32, and 40 Hz), as well as a no-vibration control, and were tested separately in three different directions (vertical, longitudinal [fore-aft], and lateral). Among these conditions, peak acceleration varied depending on frequency. Unlike active avoidance tests with electrical footshock (fear conditioning), in which the chamber is associated with the aversive stimulus, our test protocol produced no carryover effect (i.e., the avoidance result in a previous trial did not affect behavior in the following trial). In a separate experiment using vertical vibration, the peak-to-peak acceleration was fixed at 5.0532 m/s², and the displacement amplitude was varied across different frequencies (5.1, 6, 16, and 32 Hz) with corresponding amplitudes of 9.84, 7.11, 1.00, and 0.25 mm, respectively (Fig. 5D). Between trials, the chambers were cleaned using 70% ethanol and immersed in a diluted chlorine bleach solution for at least 30 minutes to eliminate residual olfactory cues.

Mouse behavior was recorded at 29.97 fps (640 × 480 pixels; VGA resolution) using a digital camera (EX-ZR1000, CASIO, Tokyo, Japan) placed above the chambers (Fig. 5A). For image analysis, the video frame rate was downsampled to ∼5 fps (one frame every 200 ms), and the analysis area was limited to the two chambers. A custom Python (version 3.8) program written by NH was used to analyze mouse position. The mouse body was detected as a binarized object (i.e., a chunk of pixels) using HSV color detection (with black representing the mouse body), followed by morphological filtering (opening and closing operations using a 4×4 kernel) implemented with OpenCV (version 4.10). The mouse position was defined as the centroid of the detected object in each frame (Fig. 5A, asterisks), and detection accuracy was confirmed by visual inspection. The percentage of time spent in the vibrator area was calculated using frame interval and the number of frames in which the centroid position was located in either “vibrator area” or “static area”. Mouse positional probabilities in the two chambers during the preference test were visualized as heatmaps of bivariate kernel density estimation (KDE) plots using a Python data visualization library, Seaborn (version 0.10.1).

### Statistical analysis and use of a Large Language Model

Statistical significance was assessed with the Mann-Whitney U-test for comparisons between awake and anesthetized mice (Figs. 1–3 and Supplementary Fig. S4; Holm-Sidak correction) and Dunn’s multiple comparison test for evaluating vibration preference relative to each corresponding control (Fig. 5), using GraphPad Prism 11 software (GraphPad, La Jolla, CA). Differences were considered statistically significant at *P <* 0.05. Pooled data are presented as mean ± SEM. ChatGPT (OpenAI, USA) and Claude (Sonnet 4.5; Anthropic, USA) were used to assist with programming for data analysis, grammatical editing and language refinement after the manuscript was drafted.

## Acknowledgements

We thank Yumi Fukuzaki for assistance with preliminary experiments and Sakura Shimakata for help with behavioral experiments. We are also grateful to Mamoru Sawada for invaluable comments and discussions.

## Author contributions

N.H. conceived the project, and M. Suzuki and N.H. designed and performed the experiments. M. Suzuki and N.H. analyzed and interpreted the data with help from M. Saito, T.U., H.I., K.S., and H.H. M. Suzuki and N.H. drafted the manuscript, and N.H. completed the paper with inputs from all the authors.

## Materials & Correspondence

Correspondence and requests for materials should be addressed to Hirokazu Hirai or Nobutake Hosoi.

## Supplementary Figures

**Supplementary Figure S1.**
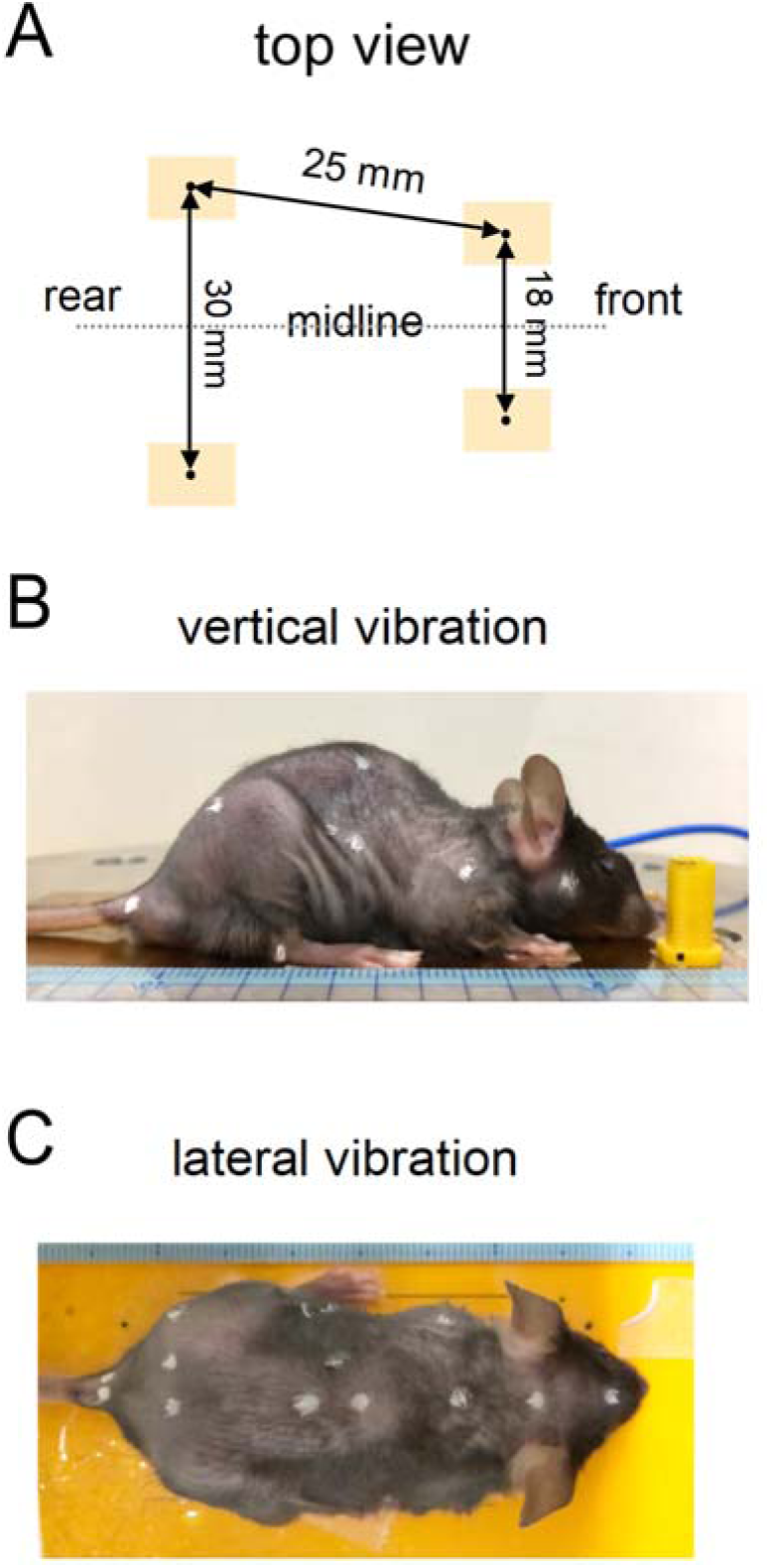
Illustrations of paw positioning in anesthetized mice to approximate a hunched posture. (A) Layout of four double-sided adhesive strips (orange rectangles; centers indicated by filled circles) used to affix all paws of the anesthetized mouse. This setup enabled the limbs to be folded under the body and kept the mouse fixed to the vibrating floor. (B and C) Illustrations including rulers with 1-mm scales and examples of mice affixed to the vibrating floor. In the vertical vibration condition (B), a yellow object with a black-marked dot at its base was taped to the floor as a reference marker for stimulation. In the lateral vibration condition (C), black-marked dots on the floor near the ‘face’ and ‘tail base’ were used as reference points for vibration.

**Supplementary Figure S2.**
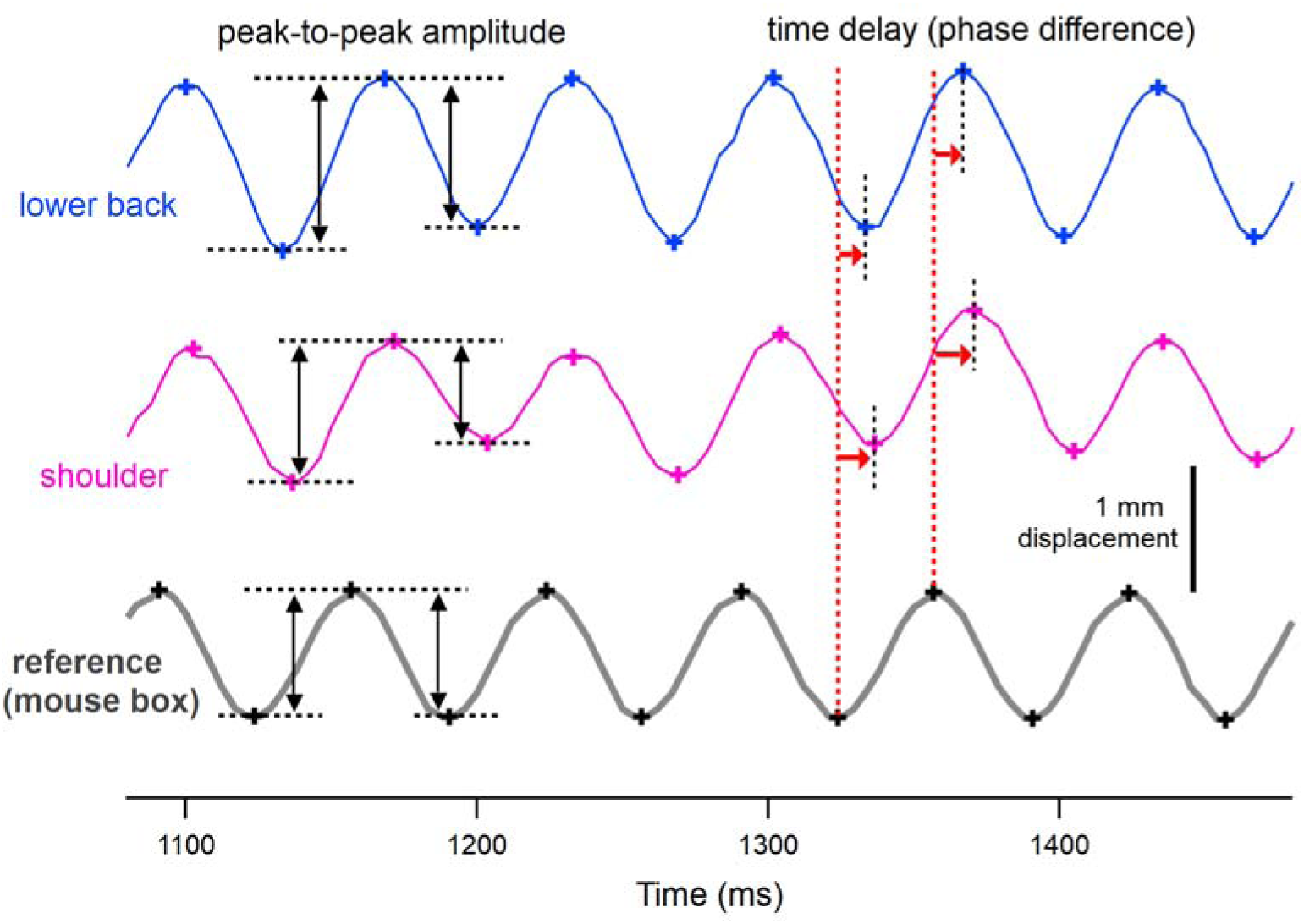
Method for measuring peak-to-peak amplitude and time delay (phase difference) from tracked displacement of mouse body parts. Representative displacement waveforms of the ‘lower back’, ‘shoulder’, and ‘reference (mouse box)’ during vertical WB vibration (sinusoidal, 15 Hz, 1 mm amplitude) in an awake mouse are shown to illustrate how amplitude and time delay (phase difference) were measured from the displacement waveforms. Notably, compared to the ‘reference (mouse box)’, the displacement of the ‘lower back’ was larger (i.e., resonance), while that of the ‘shoulder’ was slightly smaller under this experimental condition. For measurement, a stable period of the displacement waveforms was selected so that the waveforms contained at least six consecutive vibration cycles without postural change. Cross symbols indicate the detected positive and negative peak locations, which were used to measure peak-to-peak amplitude (double amplitude; black double-headed arrows) and time delay (red arrows) (see Methods). Within the stable period, multiple measurements of peak-to-peak amplitude were made, and multiple time delays were obtained based on time points of the reference peaks (vertical red dotted lines) and the corresponding peaks of each body part (vertical black dotted lines). These values (more than seven values for each amplitude and delay) were averaged to represent individual data points. Averaged time delays were converted to angular values based on the cycle time of the vibration frequency for further analysis.

**Supplementary Figure S3.**
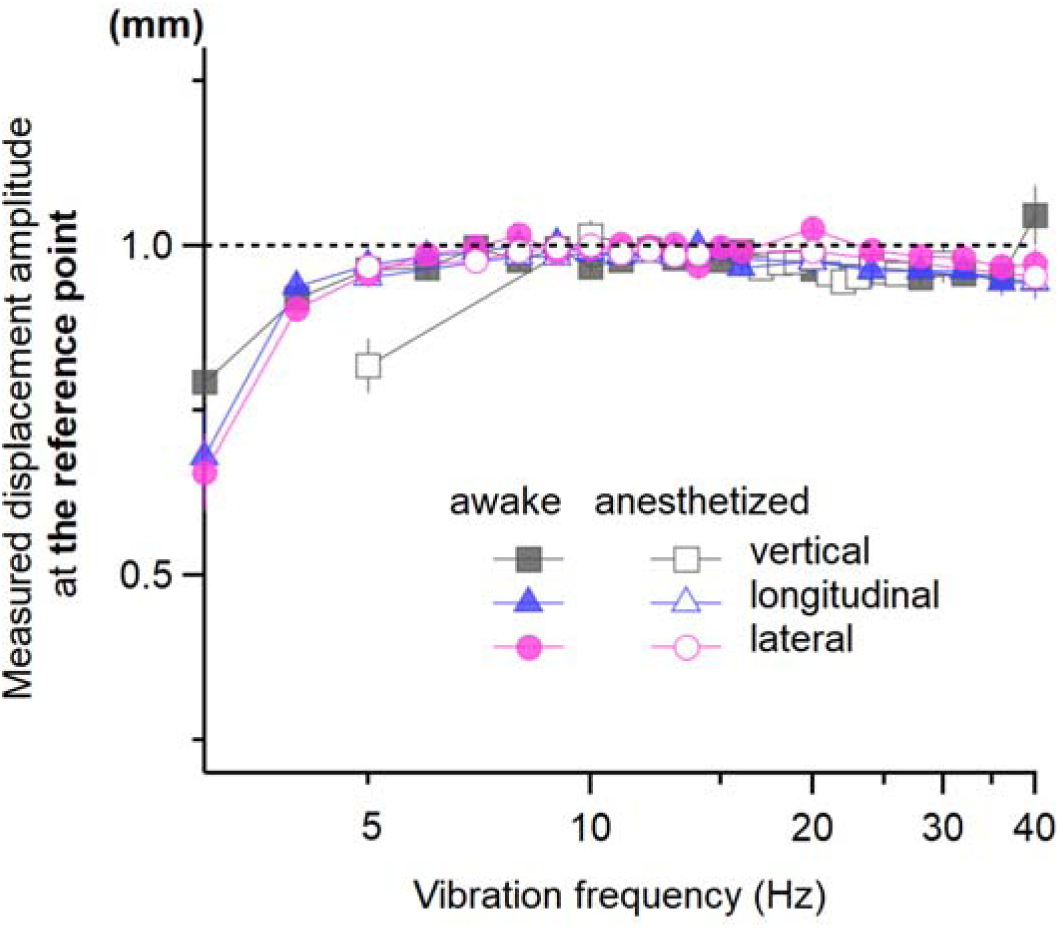
Displacement amplitudes at the reference points under various vibration conditions with different frequencies. The vibrator was configured to generate vibration stimuli with a peak-to-peak amplitude of 1 mm. To verify the accuracy of the generated vibration stimuli, actual displacement amplitudes were measured at reference points marked on the frame of the mouse chamber (see Figs. 1A–3A), on yellow reference objects affixed to the vibration platform (Supplementary Fig. S1B), or directly on the vibration platform itself (Supplementary Fig. S1C). At frequencies above 5 Hz, the measured amplitudes closely approached the intended 1-mm level (indicated by the dashed line), indicating that the vibration stimuli were well controlled in this range, regardless of vibration axis. In contrast, at lower frequencies (3–5 Hz), the displacement amplitudes fell below 1 mm due to technical limitations of the vibrator system.

**Supplementary Figure S4.**
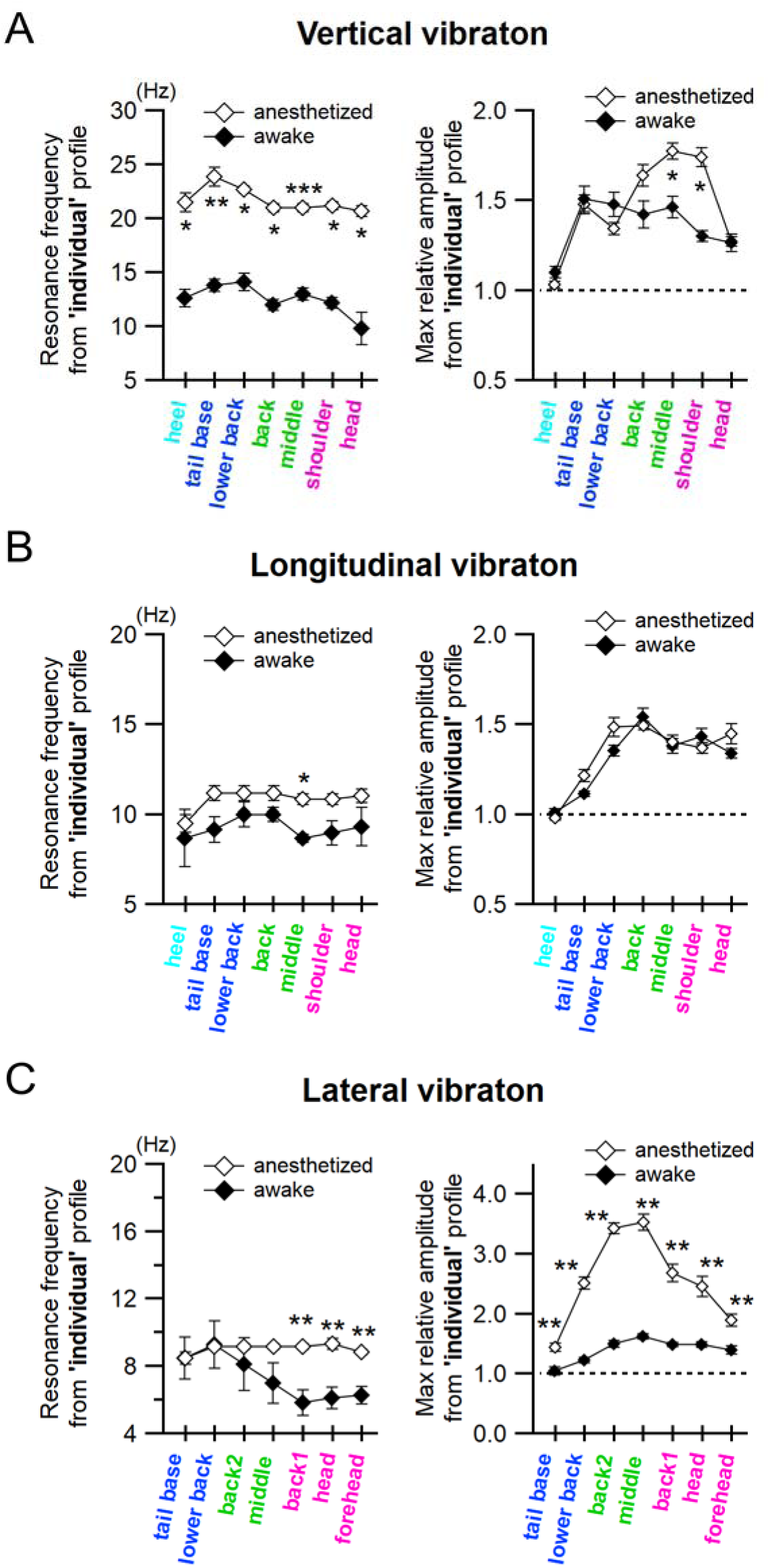
Resonance frequencies and maximum relative displacement amplitudes derived from individual mice. (A–C) In contrast to the group-averaged profiles shown in Figs. 1–3, resonance frequencies and maximum relative displacement amplitudes were obtained from individual frequency–amplitude profiles at each body part of individual mice. These values were then averaged across animals, and the mean values were plotted against each body part (left panel, resonance frequency; right panel, maximum relative displacement amplitude). (A), (B), and (C) correspond to vertical, longitudinal, and lateral WB vibrations, respectively. \*\*\**P* < 0.005, \*\**P* < 0.01, and \**P* < 0.05 (multiple Mann-Whitney tests between awake and anesthetized mice).

**Supplementary Figure S5.**
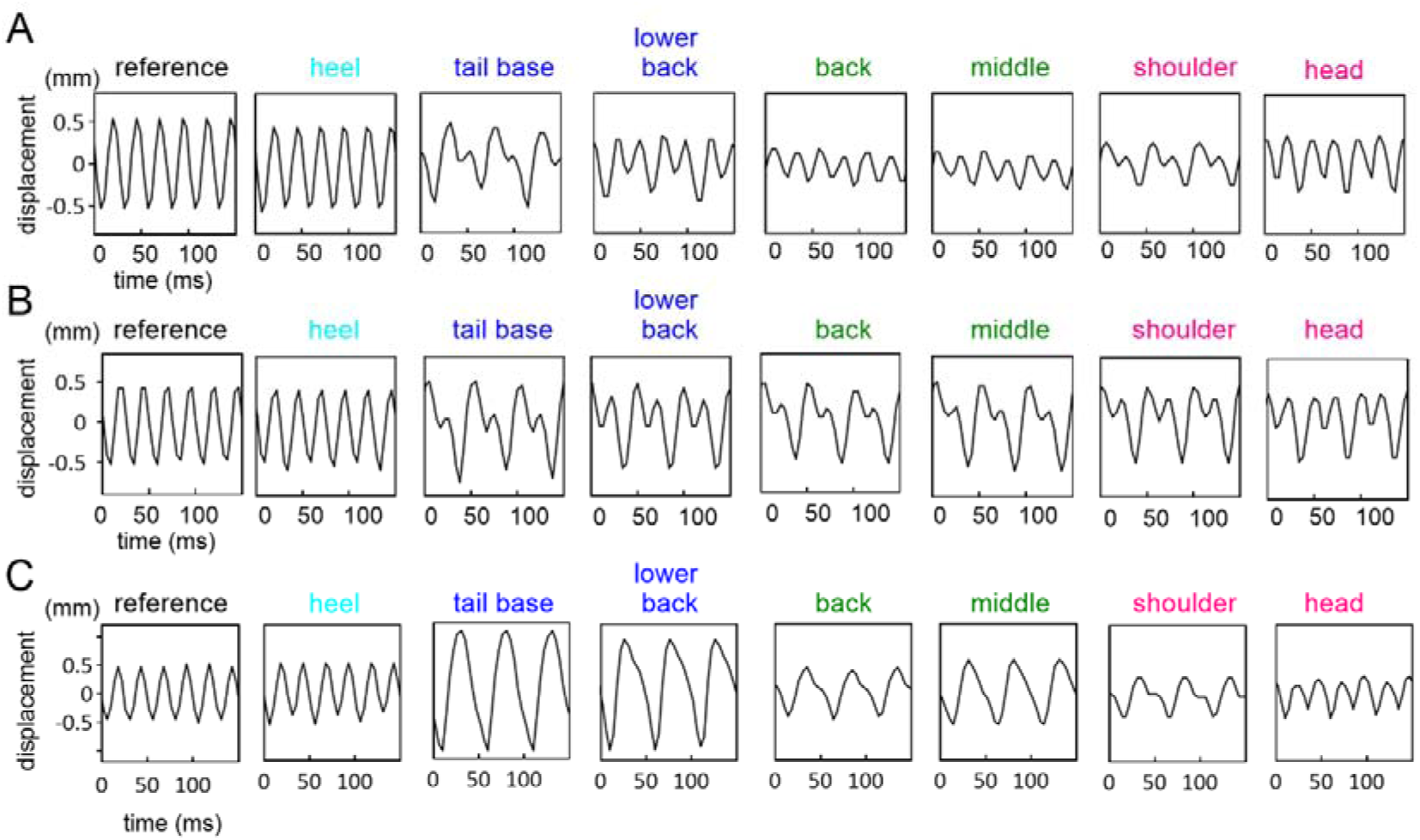
Displacement time courses at various body parts in anesthetized mice during vertical WB vibration at 40 Hz. (A–C) Displacement along the vertical vibration axis at each body part (Fig. 1A) was monitored over time in anesthetized mice exposed to vertical WB vibration at 40 Hz (sinusoidal, 1-mm peak-to-peak amplitude) using video tracking (see Methods). Panels (A), (B), and (C) show data from three different mice. The y-axis scale of the leftmost ‘reference’ graph corresponds to the scales used for the other body parts within each panel. In contrast to the ‘reference’, the displacement waveforms in all body parts except for the ‘heel’, exhibited complex patterns with pronounced positive-negative asymmetry and large inter-individual variability, making it difficult to reliably measure displacement amplitude and phase difference (see Methods). In one typical pattern of the complex displacement waveform, a relatively small positive-going hump appeared during the downward phase of displacement, as seen at the ‘tail base’ and ‘shoulder’ in (A) and (B). In another typical pattern seen at the ‘tail base’, ‘lower back’, and ‘middle’ in (C), two positive peaks appeared to merge, resulting in a broader and larger displacement waveform with a doubled apparent cycle time (∼50 ms) compared to the 25-ms cycle time of the ‘reference’ trace.

**Supplementary Figure S6.**
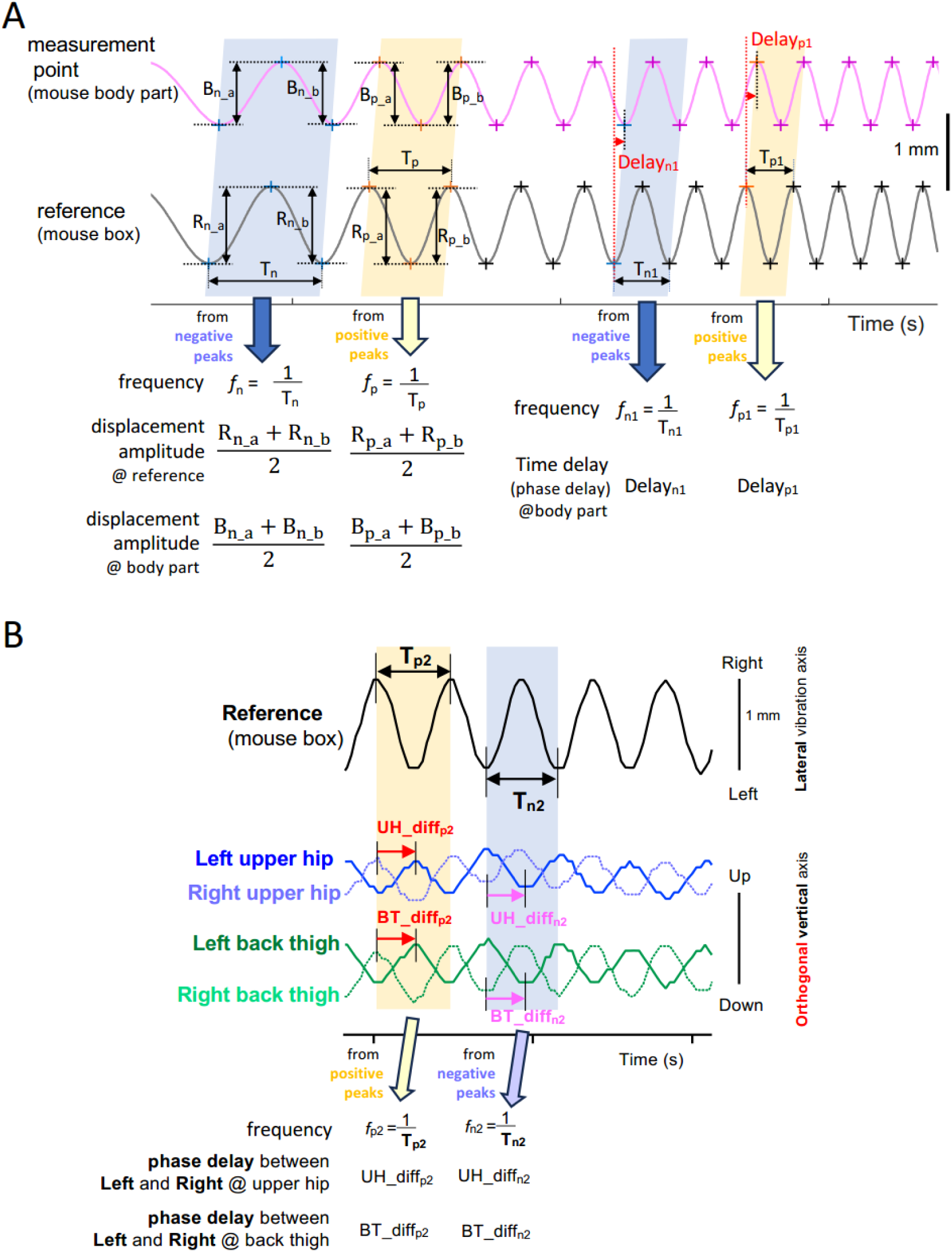
Method for measuring peak-to-peak displacement amplitude and phase delay of posterior body parts using a slowly frequency-modulated single-sweep lateral (left-right) vibration stimulus in awake mice. (A) Schematic illustration of two ideal displacement waveforms (reference and body part) in response to a frequency-modulated lateral vibration. For illustrative clarity, the frequency modulation is shown at a 14 times faster rate than the actual rate used in the experiments. Positive and negative peaks (cross symbols) were detected from all the waveforms. Time windows between two neighboring peaks with the same polarity were defined as negative-peak windows (light blue band) and positive-peak windows (cream band). For each time window, the vibration frequency was calculated from the reference trace as the inverse of the interval between successive peaks of the same polarity (T= and T=). Peak-to-peak displacement amplitudes of each trace were measured within each window, averaged, and associated with the corresponding frequency (left side). For phase delay analysis (right side), the time difference between the peak in the reference and the one in body-part trace (red arrowheads) was measured, converted to angular phase delay, and assigned to the measured frequency in each time window. Data were grouped into 0.25 Hz bins, and the mean was calculated for each bin. (B) Five representative displacement traces are shown: one reference trace (measured along the lateral axis) used to determine vibration frequency, and four body part traces (left/right ‘upper hip’ and ‘back thigh’) measured along the vertical axis. Phase delay between left and right body parts was determined by measuring the time difference between corresponding peaks (the same polarity) of left and right traces in each time window (red and magenta arrows in cream and light blue bands, respectively), assigning the result to the calculated frequency. Data were binned and averaged as described in (A).

**Supplementary Figure S7.**
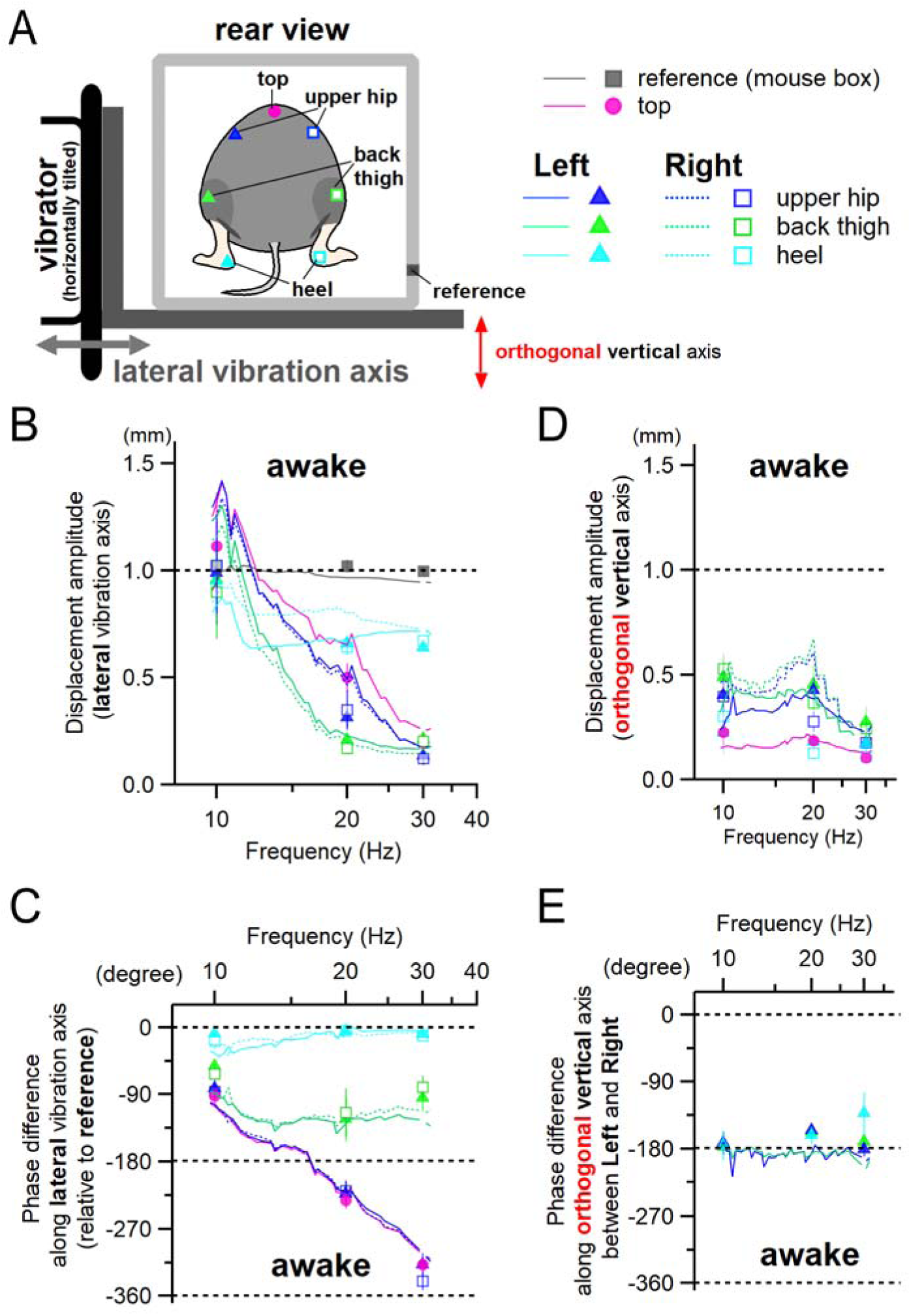
Frequency-modulated vibration stimulus reveals progressive development of phase differences exceeding -180° and antiphase movements between left and right body parts during lateral WB vibration in awake mice. (A) The experimental setup was similar to that in Figure 3, except that the movements of posterior body parts were recorded from behind during a frequency-modulated lateral WB vibration. A single-sweep frequency-modulated lateral vibration stimulus (10–30 Hz at 0.5 Hz/s, peak-to-peak displacement amplitude of 1 mm) was used to continuously assess frequency-dependent changes in phase difference. Symmetrical left and right body parts were labelled as measurement points to examine interlimb movement relationships. Color-coded symbols and lines correspond to specific left or right body parts and the associated analysis results in the other panels (B–E). (B–E) Solid and dotted lines indicate data obtained from the frequency-modulated vibration stimulus (see Supplementary Figure S6), and discrete symbols represent data from frequency-fixed lateral vibration stimuli (see Supplementary Figure S2). The consistency between the two data sets supports the validity of the frequency-modulated stimulus analysis. Data are presented in the same way as in Figure 3, except that displacement amplitudes were not normalized to the ‘reference’ displacement. (C) Phase differences along the lateral vibration axis gradually increased with stimulus frequency in upper body parts (‘top’ and ‘upper hip’), eventually exceeding -270°, whereas in lower body parts (‘back thigh’ and ‘heel’), phase differences remained below -180°. (D) Displacement along the vertical axis (orthogonal to the lateral stimulus axis) was substantial, indicating a multiple-degree-of-freedom response system. (E) Phase difference analysis along the vertical axis between left and right posterior body parts revealed a constant antiphase relationship (approximately -180°) across vibration frequencies, suggesting that these left and right body parts move in a rotational, seesaw-like manner during lateral WB vibration.

**Supplementary Video 1. Mouse body movements in response to vertical WB vibration in awake mice.**

Vibration parameters: sinusoidal displacement, 1-mm peak-to-peak amplitude at 16 Hz. Playback speed is ∼10 times slower than real time. Colored lines represent the movement trajectories of different body parts.

**Supplementary Video 2. Mouse body movements in response to longitudinal (fore-aft) WB vibration in awake mice.**

Vibration parameters: sinusoidal displacement, 1-mm peak-to-peak amplitude at 14 Hz. Playback speed is ∼10 times slower than real time. Colored lines represent the movement trajectories of different body parts.

